# PLEKHS1 drives PI3Ks and remodels pathway homeostasis in PTEN-null prostate

**DOI:** 10.1101/2023.05.18.541123

**Authors:** Tamara Chessa, Piotr Jung, Sabine Suire, Arqum Anwar, Karen E. Anderson, David Barneda, Anna Kielkowska, Barzan A. Sadiq, Sergio Felisbino, David Oxley, Dominik Spensberger, Anne Segonds-Pichon, Michael Wilson, Simon Walker, Hanneke Okkenhaug, Sabina Cosulich, Phillip T. Hawkins, Len R. Stephens

**Affiliations:** The Signalling programme, The Babraham Institute, Cambridge, UK CB22 3AT; Department of Structural and Functional Biology, São Paulo State University, Brazil; The Mass Spectrometry Facility, The Babraham Institute; The Gene Targeting Facility, The Babraham Institute; Bioinformatics, The Babraham Institute; The Imaging Facility, The Babraham Institute; Projects Group, Oncology R&D, AstraZeneca, Cambridge, U.K. 1; Now works at Division of Oncology and Center for Childhood Cancer Research, Children’s Hospital of Philadelphia, 501 Civic Center Blvd, Philadelphia, PA 19104, USA; Now works at IQVIA RDS Poland, Domaniewska 48, 02-672 Warsaw, Poland; Now works at ANU Gene Targeting Facility, Australian Phenomics Facility, John Curtin School of Medical Research, Australian National University, Canberra, Australia

**Keywords:** PLEKHS1, PI3K, PTEN, Src-family kinase, prostate

## Abstract

The PIP_3_/PI3K network is a central regulator of metabolism and is frequently activated in cancer, commonly by loss of the PIP_3_/PI(3,4)P_2_-phosphatase, PTEN. Despite huge investment, the drivers of the PI3K network in normal tissues and how they adapt to overactivation are unclear.

We find that in healthy mouse prostate PI3K activity is driven by RTK/IRS signalling and constrained by pathway-feedback. In the absence of PTEN, the network is dramatically remodelled. A poorly understood, YXXM and PIP_3_/PI(3,4)P_2_-binding PH domain-containing, adaptor, PLEKHS1, became the dominant activator and was required to sustain PIP_3_, AKT-phosphorylation and growth in PTEN-null prostate. This was because PLEKHS1 evaded pathway-feedback and experienced enhanced PI3K and SRC-family kinase-dependent phosphorylation of Y^258^XXM, eliciting PI3K activation. *hPLEKHS1-*mRNA and activating-Y^419^-phosphorylation of hSRC correlated with PI3K-pathway activity in human prostate cancers. We propose that in PTEN-null cells, receptor-independent, SRC-dependent tyrosine-phosphorylation of PLEKHS1 creates positive-feedback that escapes homeostasis, drives PIP_3_- signalling and supports tumour progression.

## Introduction

The PIP_3_ network is built around the ability of class I PI3Ks (phosphoinositide 3 kinases) to be activated by diverse signals and accelerate synthesis of the 2^nd^ messengers PIP3 (phosphatidylinositol (3,4,5)-trisphosphate) and its metabolite, PI(3,4)P_2_. PIP_3_ and PI(3,4)P_2_ are capable of driving further signalling *via* PIP_3_/PI(3,4)P_2_-binding effector proteins ultimately shaping many facets of cell biology and metabolism that support survival and growth (Fruman et al., 2017). PIP_3_ and PI(3,4)P_2_ can both be dephosphorylated by PTEN to terminate signalling (Maehama and Dixon, 1998; Malek et al., 2017). Many cancers harness mutations that drive the activity of the PIP_3_-network, including loss of PTEN activity (Li et al., 1997; Liu et al., 1997; Steck et al., 1997; Teng et al., 1997), mutations in PI3K subunits (Samuels et al., 2004) and in some cases mutations in other pathways that stimulate the pathway indirectly (e.g., SPOP in prostate cancer (Blattner et al., 2017)) and this has driven huge investment in understanding and targeting the pathway (Fruman et al., 2017; Vanhaesebroeck et al., 2021).

Class IA PI3Ks are heterodimers of a p85-family regulatory subunit (p85α, p85β or p55γ, that contain SH2 domains) and a p110-family catalytic subunit (p110α, p110β or p110δ). Both subunits are involved in the reception of diverse inputs. Signals from activated receptor tyrosine kinases (RTKs), in the form of tyrosine-phosphorylated YXXM motifs found in either the RTKs themselves (e.g., PDGFRβ) or closely associated adaptors (e.g., ERBB3, IRS or GAB-family) can bind to the p85-SH2 domains. In addition, small GTPases can interact with the catalytic subunits directly (Rodriguez-Viciana et al., 1994). Both types of input activate the catalytic subunit’s kinase domain through a web of allosteric interactions that can be hijacked by PI3K oncomutants (Burke, 2018). Activated small GTPases have a limited ability to activate class I PI3Ks in the absence of other stimuli (e.g., tyrosine phosphorylated peptides) *in vitro* or *in vivo* and work to synergistically augment activation by other inputs (Siempelkamp et al., 2017; Suire et al., 2012).

*SRC*-family tyrosine kinases (SFKs) contain SH2 and SH3 domains and can integrate a wide range of inputs, including non-receptor tyrosine kinases (such as FAK and the SRC-like PTK6 (Alwanian et al., 2022)) and RTKs. Activators, including H_2_O_2_-mediated sulfenylation of two exposed cysteines (Heppner et al., 2018) and PTK6, ultimately switch SFKs from a closed-inactive to an open-active, Y_419_- phosphorylated conformation (Y_419_ in human SRC, a highly conserved residue in the catalytic domain). The peptide sequence neighbouring tyrosine residues is important in defining motifs that are preferred as substrates by tyrosine kinases (Shah et al., 2018; Songyang et al., 1995), however, a major determinant of whether a SFK substrate is actually phosphorylated *in vivo* is co-localisation with the relevant kinase *via* a combination of SFK-SH3, and/or -SH2 domain interactions (Miller, 2003; Shvartsman et al., 2007; Songyang et al., 1995). SFKs can phosphorylate a wide range of diverse cellular substrates with important roles in, for example, regulation of the cytoskeleton, cell junctions and wound-healing (Reynolds et al., 2014). Over expression of SFKs in mouse prostate has been shown to drive tumourigenesis although the molecular targets of the SFKs that mediate this are unclear (Cai et al., 2011). SFKs have also been suggested to be key upstream coordinators of many of the changes that occur in the phospho-tyrosine landscape during prostate tumour progression (Drake et al., 2013; Drake et al., 2016) and to drive EMT, invasion and metastasis (Ortiz et al., 2021). Very few SFK substrates are phosphorylated within YXXM motifs and in most cases SFKs activate PI3Ks indirectly *via* other kinases. A small subset of non-YXXM, potential SFK-substrate sites have, however, been suggested to bind p85s and activate PI3K signalling (Moon et al., 2005; Riggins et al., 2003).

Use of PI3K-pathway inhibitors in cancers, or transformed cell lines, with mutations leading to elevated PI3K-network activity has revealed that there is powerful feedback from active mTORC1. Firstly, *via* p70_S6K_ and/or Grb10 which leads to suppression of the expression (at both mRNA and protein levels) and phosphorylation of key RTKs (e.g., IGFR1, EGFR) and their adaptors (e.g., IRS). Secondly, via 4E-BP1 that leads to the increased translation and accumulation of PTEN. This feedback acts homeostatically, but against therapy, to return network activity towards its set-point (Chakrabarty et al., 2012; Chandarlapaty et al., 2011; Hsu et al., 2011; Mukherjee et al., 2021; Rodrik-Outmezguine et al., 2011).

Drivers of the PI3K pathway in cancer and cancer cell lines have been identified by antibody-directed p85-pulldowns followed by immuno-blotting for candidates. These studies repeatedly identified molecules like ERBB3, GAB1/2, PDGFRβ and IRS-1 proteins as key (Ebi et al., 2011; Engelman et al., 2005; Engelman et al., 2007; Stommel et al., 2007; Turke et al., 2010) (in one case using proteomics to come to a similar conclusion (Yang et al., 2011)). The primary physiological molecular activators of class IA PI3Ks, however, remain poorly understood, consequently, important questions about whether or how physiological PI3K networks adapt or remodel in the face of sustained perturbations also remains unclear.

We set out to systematically search for direct activators of the PI3K pathway in both normal mouse prostate and also prostate in which the pathway had been chronically activated by loss of PTEN. This mouse model, in which PTEN can be conditionally ablated from prostate epithelial cells, has been characterised extensively (*Pten^loxP/loxP^ x PbCre4*^+^)(Trotman et al., 2003; Wu et al., 2001) and shown to involve a very substantial increase in the levels of both PI(3,4)P_2_ and PIP_3_ (Malek et al., 2017). The protein phosphatase activity of PTEN has been demonstrated to dephosphorylate the Src-like tyrosine kinase PTK6 at the activating Y^342^ site (Wozniak et al., 2017). Hence loss of PTEN can lead to activation of PTK6 and as a result SRC (Alwanian et al., 2022). Young mice display prostate hyperplasia progressing to low and then high-grade PIN from 3–7 months with invasive adenocarcinoma emerging from 12 months (Jurmeister et al., 2018).

We have previously generated mice in which their endogenous p85α or β proteins were fully biotinylated on 17aa C-terminal avi-tags (Beckett et al., 1999; Tsolakos et al., 2018) (the mice also expressed a “mammalised” (m) version of a prokaryotic biotin ligase BirA, that was knocked-into, and widely expressed from, the *ROSA26* locus and capable of biotinylating Avi-tags *in vivo*; e.g., *p85α^Avi/Avi^* x *mBirA*^+/-^). We have shown that p85α and β from these mice are expressed at normal levels and can be very efficiently recovered from tissue lysates in rapid streptavidin-mediated pulldowns (Tsolakos et al., 2018).

By crossing mice expressing Avi-tagged p85s and mBirA with mice in which PTEN could be conditionally deleted in the prostate epithelium we generated unique tools to define the endogenous PI3K signalling network in the context of either physiological or hyper-activated PIP_3_ signalling.

## Results

### The p85-PI3K interactome is dramatically remodelled upon loss of PTEN in mouse prostate

We aimed to define and semi-quantify the family of proteins associated with p85α and β PI3K complexes in wild-type and PTEN-KO mouse prostate by proteomic analysis of streptavidin pulldowns and use of TMT-labelling to bar-code, multiplex and compare the relative recovery of proteins between control pulldowns (no Avi-tag but expressing mBirA, e.g., *p85α^WT/WT^ x mBirA*^+/-^) and specific pulldowns (with an Avi-tag and expressing mBirA, e.g., *p85α^Avi/Avi^ x mBirA*^+/-^). The work flow is summarised in Figure 1A and S1A (note; all mice express mBirA but this is omitted from Figure 1A for clarity). The results showed the presence of Avi-tags on endogenous p85s had no effect on either their expression (Figure S1B) nor PI3K activity as read-out by PIP_3_ accumulation in either wild-type or PTEN-KO prostate (Figure S1E) and that the protocol had worked effectively with efficient pulldown and specific recovery of p85α-Avi and p85β-Avi and associated p110s (Figure 1 and S1B-D). Twenty-four proteins were identified that were significantly recovered in pulldowns from *p85α^Avi/Avi^* or *p85β^Avi/Avi^* compared to *p85^WT/WT^* healthy prostate (Figure 1B). When this data is presented in a form that enables the relative moles of the associated proteins that were recovered to be compared on a common scale (Figure S1F) then it is clear that in wild-type, healthy mouse prostate tissue IRS1 dominated this list and can be seen as the likely primary driver of the PIP_3_ network. In PTEN-null tissue the picture of proteins recovered with p85s was changed entirely. IRS proteins were much reduced with smaller but significant reductions in recovery of EGFR and SHC (Figure 1B and C); some interactors were unchanged (Figure 1B and D), whilst several increased substantially (Figure 1B and E).

**Figure 1.**
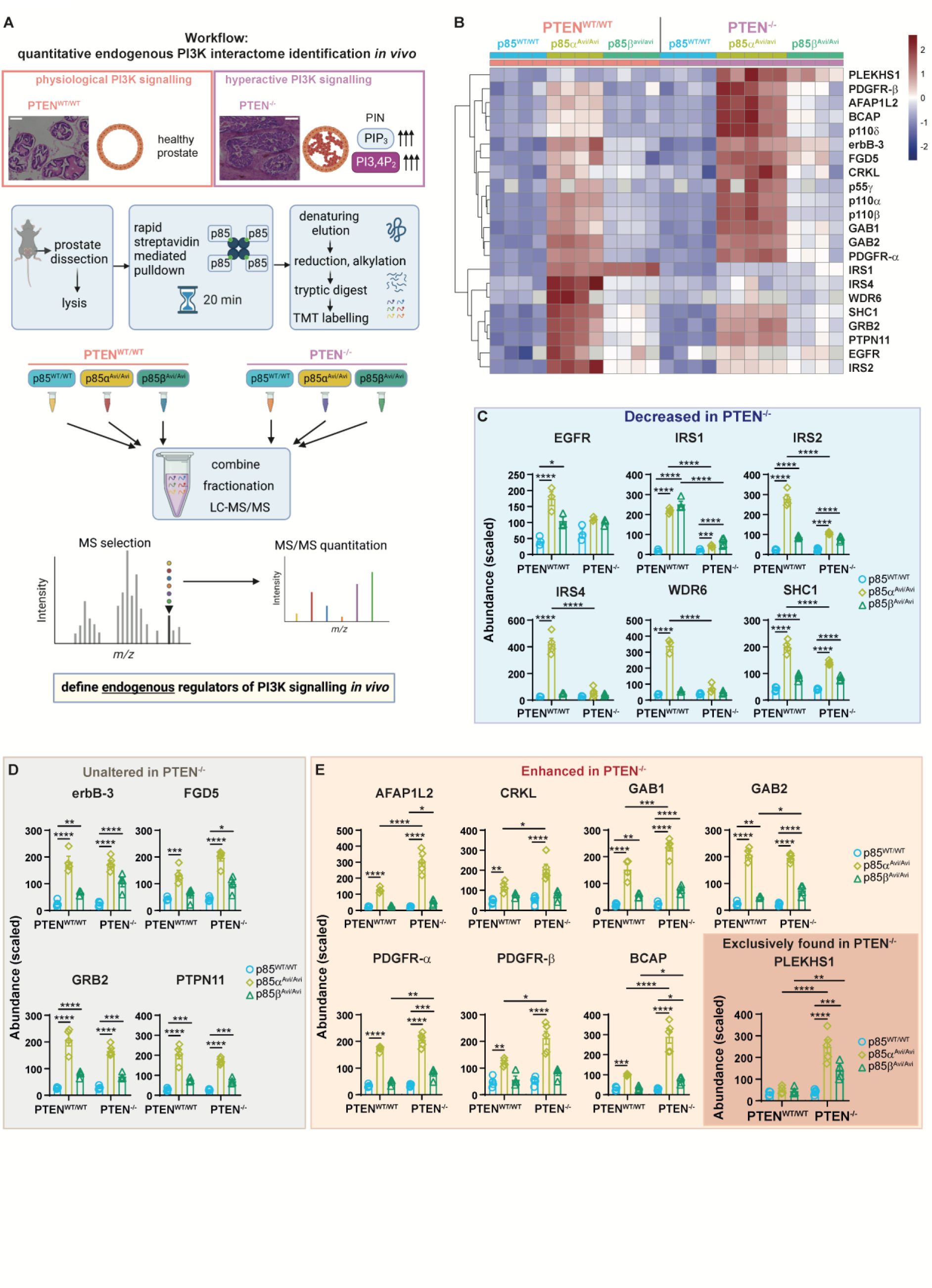
The p85 interactome is dramatically rewired in PTEN-KO prostate. **A,** Methodology for identification of p85α- and p85β-interactors in *Pten^WT/WT^* and *Pten^-/-^* prostates (12 weeks). Scale bar, 100μm. **B,** Proteins that were significantly more enriched in *p85α* or *β^Avi/Avi^* pulldowns compared to *p85^WT/WT^* pulldowns were defined as specific p85-interactors. Multiple two-sided unpaired *t*-tests (Holm correction) were performed. A stringent significance threshold (p<0.01) was applied to identify proteins with increased detection above *p85^WT/WT^* controls and specific interactors are visualised in a heatmap (clustering on z-scores of row normalised values). **C,** The data presented are scaled abundances of p85-interactors that were decreased in PTEN^-/-^ tissue (n=4-5 biological replicates per genotype, as indicated in Fig 1b and as described in the Methods and the legend to Figure S 1C and D. Data for PI3K subunits is shown in FigureS1C and D). For statistical analysis, 2-way ANOVAs were performed followed by Holm-Šídák’s multiple comparisons tests. Adjusted P value summaries <0.05 are indicated. **D**, As above, except the data are scaled abundances of p85-interactors that were unchanged in PTEN^-/-^ tissue. **E**, As above, except the data are scaled abundances of p85-interactors that were increased in PTEN^-/-^ tissue.

Ranking the YXXM-containing adaptors according to their fold increase in PTEN-null compared to wild-type prostate identified PLEKHS1 (Pleckstrin Homology Domain Containing S1), a poorly understood, GAB-related adaptor, as the most increased (Figure 1B, 1E and S 1H). PLEKHS1, and several other key interactors, were also identified in p85-pulldowns from PTEN-KO and wild-type prostate by immuno-blotting (Figure S1G; note, murine and hPLEKHS1 run anomalously during SDS-PAGE, the predicted sizes of their longest transcripts are 53-55kD but they run with a mobility indicative of 70kD, see the legend to Figure S1G).

### IRS but not PLEKHS1-, mediated pathway activation is sensitive to pathway feedback

To investigate whether any of these changed associations were dependent on acute PI3K activity, mice were dosed with a pan-class I PI3K inhibitor GDC-0941 (pictilisib (Folkes et al., 2008) Figure 2A). The inhibitor reduced PIP_3_ levels and AKT phosphorylation in the prostate, as expected (Figure S2A and C). This treatment regime had no effect on the recovery of most p85-interactors (Figure 2D, 2E and S2B), however, recovery of IRS1 and 2 from both wild-type and PTEN-KO prostate was significantly increased (Figure 2C). In direct contrast, PI3K inhibition caused a decrease in PLEKHS1 recovery with p85s (Figure 2B).

**Figure 2.**
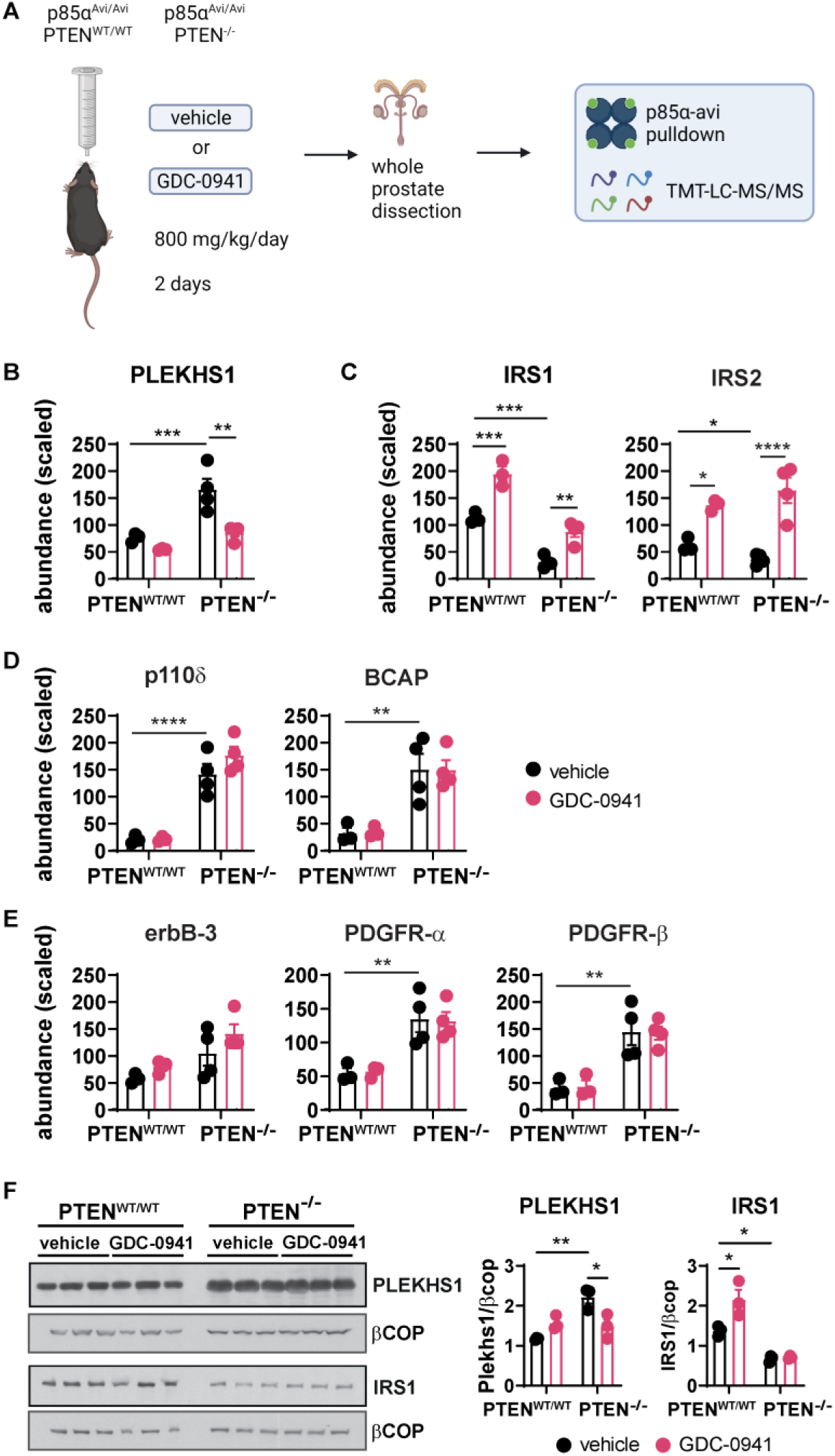
IRS but not PLEKHS1-, mediated pathway activation is sensitive to pathway feedback. **A,** Diagram of dosing schedule of p85α^Avi/Avi^ PTEN^WT/WT^ and p85α^Avi/Avi^ PTEN^-/-^ mice with pan-class I PI3K inhibitor GDC-0941 and experimental workflow. **B-E,** Targeted TMT-LC-MS/MS analysis of the indicated proteins in p85α^Avi/Avi^, PTEN^WT/WT^ and p85α^Avi/Avi^, PTEN^-/-^ prostates (24-32 weeks), treated with vehicle or GDC-0941. Data are means ± SEM for n=3-4 biological replicates per genotype/condition, run in 2 separate cohorts (p85α^Avi/Avi^, PTEN^WT/WT^: n=3; p85α^Avi/Avi^, PTEN^-/-^: n=4). For statistical analysis, 2-way ANOVAs were performed followed by Holm-Šídák’s multiple comparisons tests. **F,** PLEKHS1 (G17) and IRS1 immunoblots of total lysates from p85α^Avi/Avi^, PTEN^WT/WT^ and p85α^Avi/Avi^, PTEN^-/-^ prostates, treated with vehicle or GDC-0941 (20 μg total protein/lane). Data are means ± SEM of 3 biological replicates per genotype/condition, run as a single cohort. For statistical analysis, 2-way ANOVAs were performed followed by Holm-Šídák’s multiple comparisons tests.

To understand the origin of these changes in p85-interactomes we examined published data measuring the mRNAs of these interactors in healthy or PTEN-KO mouse prostate (NCBI GEO accession number GSE94574, comparing normal and PIN) and measured their abundance in prostate lysates. This revealed that the changes in p85-interactome upon loss of PTEN were very unlikely to be driven by changed expression with the exception of the IRS family; where the levels of their mRNA and protein were substantially and significantly reduced in PTEN-KO prostate (Figure S2A, S2C and S3A) and this was partially reversed by PI3K inhibition (Figure 2F). In contrast, there was a small decrease in *Plekhs1*-mRNA (Figure S3A) and a modest increase in PLEKHS1 protein in the absence of PTEN, with the latter reduced by PI3K inhibitors (Figure 2F).

Collectively, the results in Figures 1 and 2 are consistent with PI3K-pathway feedback being active in healthy prostate tissue leading to physiological suppression of IRS activation of PI3K signalling. In PTEN-KO prostate tissue that feedback intensified and further suppressed expression of IRS proteins (and WDR6, a known IRS4-interactor) and thus their ability to interact with p85s. The outcome is that IRS proteins become dramatically reduced and relatively minor drivers of the PI3K pathway in PTEN-KO prostate (Figure S1F).

Public data sets show that m*Plekhs1* mRNA is differentially expressed in a small number of tissues including prostate and we have confirmed that PLEKHS1 protein is expressed in mouse prostate (see below). To understand which cell types in the prostate expressed PLEKHS1 and what might underlie the increase in PLEKHS1 expression in PTEN-KO prostate, we dissociated mouse prostate tissue and sorted the resulting cell suspension into basal-epithelial, luminal-epithelial and residual (including stromal and immune cells) cell bins and measured PLEKHS1 by immuno blotting. This revealed that the large majority of PLEKHS1 was detected in epithelial cells in both wild-type and PTEN-KO prostate and was significantly enriched in luminal compared to basal cells from both genotypes (Figure S4A and B).

Importantly, the expression of PLEKHS1 was the same in luminal cells purified from PTEN-KO or wild-type prostate; suggesting that the increase in PLEKHS1 expression in PTEN-KO prostate resulted from an increase in the relative size of the luminal cell compartment in the prostate (Figure S4A and B) and not an increase in its concentration in prostate luminal cells. Collectively, these results suggest that, in contrast to IRS proteins, PLEKHS1 is not subject to pathway feedback and that its increased association with p85s in PTEN-KO tissue is not a result of a dramatic change in its expression.

### A subset of p85-interactors that are increased in PTEN-null prostate are a result of immune infiltration

Of the group of interactors that were increased in PTEN-KO prostate, two, BCAP/PIK3AP and p110δ are recognised as differentially expressed in murine immune cells, particularly macrophages (Figure 1B, S3B and S3C). Furthermore, the recovery of CSF1R in p85-Avi pulldowns was greater from PTEN-null compared to control prostate, although this data was not statistically significant because CSF1R peptides were detected in only 2/5 samples (Figure S3B). This may reflect infiltration of myeloid-derived suppressor cells, with active CSF1R/BCAP/PI3Kδ signalling, into the hyperplasic tissue. This would be consistent with the activated immune phenotype of the *Pten^loxP/loxP^ x PBCre4*^+^ model (Zhang et al., 2020a). Interestingly, the recovery of this group of interactors with p85s was unchanged by inhibition of PI3K activity (Figure 2D) suggesting neither the assembly of these signalling complexes nor the infiltration of the relevant immune cells into the PTEN-KO prostate was acutely dependent on PI3K activity.

### PLEKHS1 can be phosphorylated by Src-family kinases on Y^258^XXM and bind and activate PI3K signalling and contains a PH domain that can bind PI(3,4)P_2_ and PIP_3_

Previous work has shown that the YXXM motif in hPLEKHS1 (equivalent to Y^258^ in the mouse, Figure 3A) can be tyrosine phosphorylated by a variety of non-receptor tyrosine kinases and bind to the C-terminal SH2 domain of p55γ in heterologous expression systems (Grossmann et al., 2015). Furthering this work, we used Hela and LNCaP cells (a human, PTEN-null, prostate cancer cell line) to demonstrate that heterologous Y^527^F-*cSrc* but not K^295^F-*cSrc* (constitutively active and inactive alleles, respectively, Figure 3B) drives both phosphorylation of Y^258^ in heterologous mPLEKHS1 and Y^258^-mPLEKHS1-dependent binding of mPLEKHS1 to endogenous p55γ and p85 subunits. In LNCaP cells, over-expression of mPLEKHS1 alone increased PIP_3_ levels in a manner disrupted by Y^258^F mutation of PLEKHS1, suggesting that endogenous kinases had phosphorylated Y^258^-PLEKHS1 and driven activation of class IA PI3K signalling (Figure 3C).

**Figure 3.**
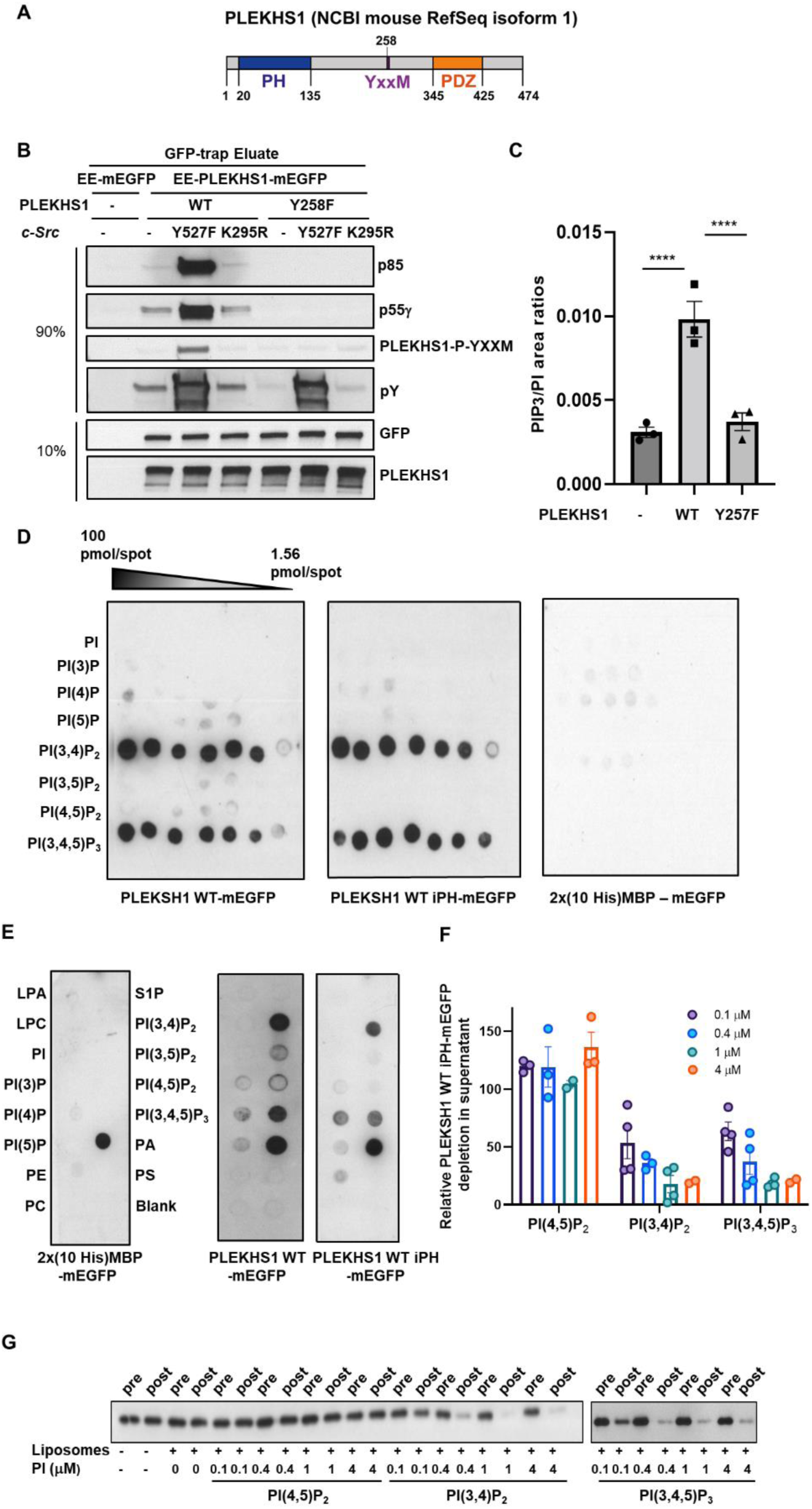
PLEKHS1 can be phosphorylated by Src-family kinases on Y^258^XXM and bind and activate PI3K signalling and contains a PH domain that can bind PI(3,4)P_2_ and PIP_3_. **A,** Domain structure of the canonical sequence of mouse PLEKHS1 (NCBI RefSeq isoform 1). **B,** LNCaP cells were transfected with tagged *mPlekhs1* (WT-EE-*Plekhs1*-mEGFP or Y^258^F-EE-*Plekhs1*-mEGFP) or vector (EE-mEGFP) alone, with or without constitutively active *cSrc* (Y527F) or kinase-dead *cSrc* (K295R). Lysates were prepared, subjected to GFP-trap pulldown and the eluates were immuno-blotted as indicated to assay PLEKHS1 interaction with PI3K regulatory subunits (p85, p55γ) and PLEKHS1 Tyr phosphorylation (total or pY^258^xxM) with GFP and PLEKHS1 as input-controls. The immuno-blots are representative of 2 biological replicates. **C,** LNCaP cells were transfected with WT-EE-*mPlekhs1*-mEGFP (WT), Y258F-EE-*mPlekhs1*-mEGFP (Y258F) or vector alone (EV). The cells were extracted and total PIP^3^ was quantified and normalised to total PI measured in the same sample. The data are presented as means ± SEM of 3 biological replicates, significance was tested with a One-way ANOVA followed by Holm-Šídák’s multiple comparisons tests. **D and E,** PIP arrays (**D**) and strips (**E**) were incubated with GFP-conjugated probes; PLEKHS1 WT (10-459), its isolated PH domain (10-134) or a control protein (the protein generated by the expression vector without a PLEKHS1 insert; 2x(10His) MBP-mEGFP). The probes were visualised with anti-GFP primary and HRP conjugated anti-mouse secondary antibodies. **D,** The PIP arrays were spotted with a range of amounts (1.56-100 pmol) of eight synthetic phosphoinositides. **F-G,** Sucrose-loaded vesicles containing a mix of PS, PE, PC, sphingomyelin and a range of concentrations of PIP_3_ or PIP_2_s (0.1 μM to 4 μM or 0.026 to 1 mol%) were incubated with GFP-conjugated iPH PLEKHS1. The assays were ultracentrifuged to sediment the liposomes and aliquots from before and after sedimentation were resolved by SDS-PAGE and immuno-blotted with an anti-GFP primary and HRP conjugated anti-mouse secondary antibodies to determine the proportion of iPH that remained in the supernatant (**G**). Data was quantified (as a % of input remaining in the supernatant) from a minimum of 3 independent experiments (all the independent replicates are plotted) and are presented as means ± SEM **(F)**.

PLEKHS1 contains a PH domain, with key similarities to PIP_3_-binding PH domains (Figure 3A). We tested whether full-length, purified, bacterially-expressed PLEKHS1 or its isolated PH domain could selectively bind phosphoinositides presented in either protein-lipid overlay assays or sucrose-loaded liposomes with a plasma membrane-like mix of phospholipids. Both constructs selectively bound PIP_3_ and PI(3,4)P_2_ with similar apparent affinity (Figure 3D-G).

### Pull-down of endogenous PLEKHS1 shows interactions with PI3Ks and 14:3:3 proteins are increased in PTEN-null tissue

To independently test the idea that PLEKHS1 can interact with p85s and that this is augmented in PTEN-null prostate and to improve our ability to quantify PLEKHS1 and its phosphorylation *in vivo* we generated mouse strains in which an Avi-tag was knocked-in to the 3’-end of the endogenous *Plekhs1* locus (Figure S4C).

*Plekhs1*^Avi/Avi^ mice were viable with no overt, or prostate-growth/morphology, phenotypes. Expression of PLEKHS1was unchanged in *Plekhs1*^Avi/Avi^ prostate and PLEKHS1-Avi could be completely, specifically depleted from lysates of m*BirA*^+/-^ mice with streptavidin-beads (Figure S4D). We assessed the distribution of PLEKHS1 between different tissues in PLEKHS1-Avi expressing mice by immuno blotting with both anti-PLEKHS1 and anti-Avi antibodies (Figure S4E). Those experiments revealed that PLEKHS1 is differentially expressed with relatively high levels in prostate, oviduct and uterus and undetectable levels in the other tissues we sampled; broadly confirming the published tissue distribution of *mPlekhs1* mRNAs. *Plekhs1*^Avi/Avi^ mice were interbred with *mBirA^+/-^*, *Pten^loxP/loxP^* and *PbCre4*^+^ mice to yield the relevant genotypes for experiments. Streptavidin-pulldowns from wild-type or *Pten*^-/-^ prostate revealed a dramatic increase in association of PLEKHS1-Avi with class IA PI3K subunits in PTEN-KO tissue (Figure 4B), confirming the above results with p85α-Avi and p85β-Avi-expressing prostate. These experiments also found a large increase in recovery of 14:3:3 proteins with PLEKHS1-Avi from PTEN-KO prostate (Figure 4C). No tyrosine kinases, that might phosphorylate Y^258^-PLEKHS1, were identified as significant interactors. String and gene ontology analysis of PLEKHS1-interacting proteins whose recovery was significantly increased from PTEN-null compared to wild-type prostate did not implicate RTKs or their proximal adaptors (Figure 4A), unlike similar analyses of public, experimental data for mGAB or mIRS proteins (Figure 4E and F). These data suggest that in PTEN-null tissue PLEKHS1 is, at most, weakly regulated by RTKs.

**Figure 4.**
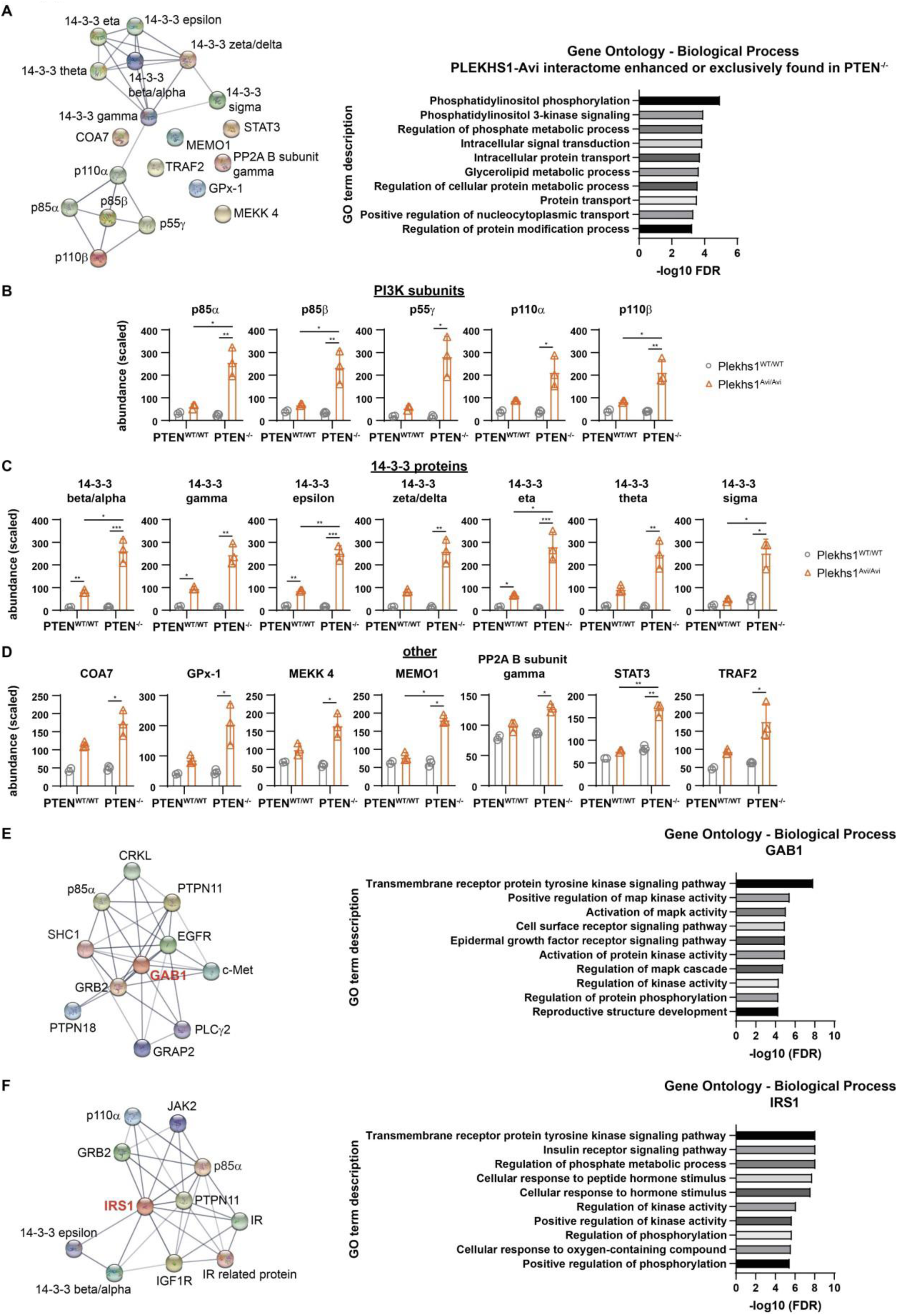
Pull-down of endogenous PLEKHS1 shows interactions with PI3Ks and 14:3:3 proteins are increased in PTEN-null tissue. **A,** STRING network analysis (experimentally determined interactions only) of specific PLEKHS1-Avi interactors (proteins recovered in significantly greater amounts in pulldowns from *Plekhs1^Avi/Avi^* x *mBirA^+/-^* compared to *Plekhs1^WT/WT^* x *mBirA^+/-^* tissue) that were recovered in significantly greater abundance from *Pten^-/-^* prostate, or exclusively found in *Pten^-/-^* prostate (12-15 weeks). Network nodes represent proteins; edges represent protein-protein associations, including functional and physical interactions. Line thickness indicates the strength of data support. **B, C and D** Scaled abundances of PI3K subunits (**B**) and 14-3-3 proteins (**C**) and other interactors (**D**) showing significant enrichment in *Plekhs1^Avi/Avi^ x Pten^WT/WT^* and/or *Plekhs1^Avi/Avi^ x Pten^-/-^* prostates (12-15 weeks) above their no-Avi controls (*Plekhs1^WT/WT^ x Pten^WT/WT^* and/or *Plekhs1^WT/WT^ x Pten^-/-^*) as calculated in Proteome Discoverer software. The experiment was run as a single cohort of n=3 biological replicates per genotype, except *Plekhs1^WT/WT^ x Pten^-/-^* where n=2. For statistical analysis, a 2-way ANOVA was performed with adjusted P values calculated using the Benjamini-Hochberg method. Adjusted P value summaries <0.05 are indicated. Data are means ± SEM or means ± range (n=2). **E and F**, STRING network analysis (experimentally determined interactions only) of mouse GAB1 (**E**) and mouse IRS1 **(F**). A maximum number of 10 interactions is shown. Network nodes represent proteins; edges represent protein-protein associations, including functional and physical interactions. Line thickness indicates the strength of data support.

### Y^258^XXM-PLEKHS1 is phosphorylated in a PI3K and SRC-family kinase dependent fashion

We investigated whether phosphorylation of Y^258^-mPLEKHS1 could be detected *in vivo* and if it was changed upon loss of PTEN and/or inhibition of PI3K activity. Phospho-proteomic analysis of PLEKHS1-Avi-pulldowns detected phosphorylation of Y^258^-PLEKHS1, that was significantly increased in PTEN-KO prostate, and a further fifteen serine and five threonine residues that were unambiguously phosphorylated *in vivo* (Figure S5A). Pulldowns from lysates of wild-type and PTEN-KO prostate with anti-phospho-tyrosine antibodies followed by immuno-blotting with anti-PLEKHS1 antibodies showed that there was an increase in total tyrosine phosphorylation of PLEKHS1 in PTEN-KO prostate (Figure S5B). Pulldowns of PLEKHS1-Avi from wild type or *Pten*^-/-^ prostate from mice either treated with GDC-0941 or its vehicle, followed by immuno-blotting with the anti-phospho-Y^258^-PLEKHS1 antibody (validated in Figure 3B) revealed that phosphorylation of Y^258^-PLEKHS1 was increased in PTEN-KO prostate and this was significantly reduced by inhibition of PI3K activity (Figure 5A). Further, the reduction in phosphorylation of Y^258^-PLEKHS1 caused by PI3K inhibition was paralleled by reductions in recovery of p85α and p55γ (Figure 5A). Taken together, these results show phosphorylation of Y^258^-PLEKHS1 is increased in PTEN-KO tissue and sensitive to PI3K inhibitors. The ability of the PH domain of PLEKHS1 to bind PIP_3_ and PI(3,4)P_2_ offers a simple molecular explanation: the dramatically increased levels of those lipids in PTEN-KO prostate led to a redistribution of PLEKHS1 that is required for phosphorylation.

**Figure 5.**
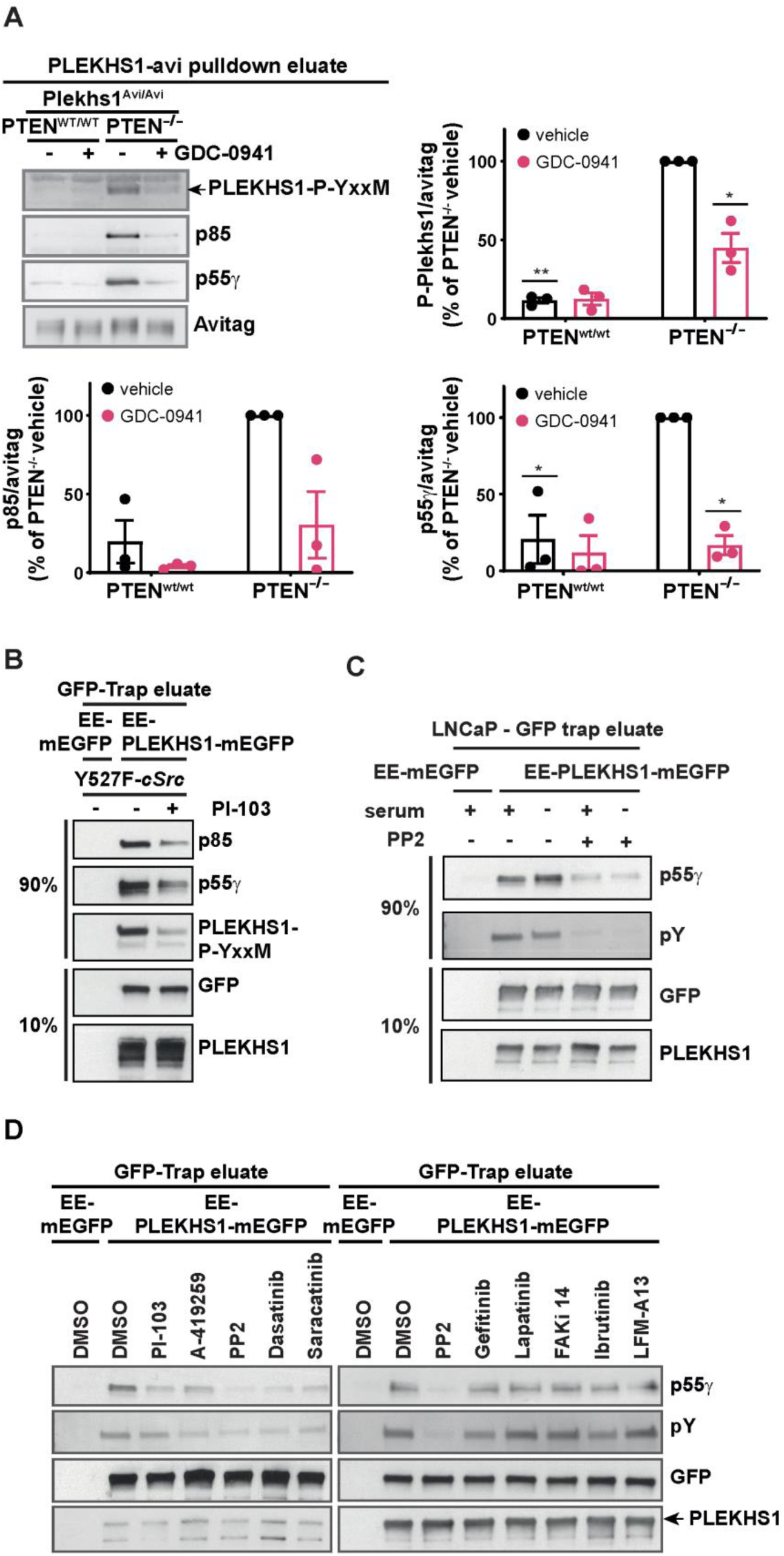
Y^258^XXM-PLEKHS1 is phosphorylated in a PI3K and SRC-family kinase dependent fashion. **A**, PLEKHS1-Avi was pulled-down from prostates lysates from *Pten^WT/WT^* or *Pten^-/-^* mice (all expressing *mBirA^+/-^, 12-15w)* that had been treated with either vehicle or PI3Ki (GDC-0941, 800 mg/kg/day for 2 days). Phosphorylation of PLEKHS1 on Y^258^xxM and recovery of total p85 and p55γ were quantified by immuno-blotting as indicated. Data were obtained from 3 independent experiments (in each experiment a minimum of 3 prostates were pooled from *PTEN^WT/WT^* mice and a minimum of 2 prostates were pooled from *PTEN^-/-^* mice) and are presented in the form of %s of the vehicle-treated, *Plekhs1^Avi/Avi^ x Pten^-/-^* samples and are means ± SEM. For statistical analysis, multiple two-tailed, one-sample T-tests were performed on baseline corrected data (all compared to the data defining 100%) using the Holm-Šídák method. Adjusted P value summaries <0.05 are indicated. **B,** LNCaP cells were transfected with tagged *mPlekhs1* (WT-EE-Plekhs1-mEGFP) or vector (EE-mEGFP) alone, with or without constitutively active *cSrc* (Y527F) or pre-treatment with various inhibitors or their vehicle as indicated. Lysates were prepared, subjected to GFP-trap pulldown and the eluates were immuno-blotted as indicated. Cells were pre-treated with PI3Ki (2 mM PI-103) or vehicle (0.1% DMSO) for 1hr, eluates were immunoblotted to assay PLEKHS1 interaction with PI3K regulatory subunits (p85, p55γ), phosphorylation of Y^258^XXM-PLEKHS1 with GFP and PLEKHS1 as input controls (n=1). **C,** GFP-trap pulldown from LNCaP cells following transfection of tagged mPLEKHS1 (EE-PLEKHS1-mEGFP). Empty vector (EE-mEGFP) included as negative control. Cells were serum starved (16 hr) and pre-treated with either vehicle DMSO (0.1 %) or SFK-inhibitor PP2 (10 μM). Eluates were immunoblotted to assay PLEKHS1 interaction with p55γ and PLEKHS1 total Tyr phosphorylation (n=2). **D**, Cells were pre-treated. with vehicle (0.1% DMSO), PI3Ki (2 mM PI-103), Src-family kinase inhibitors A419259 (1 µM), PP2 (10 µM), Dasatinib (100 nM), Saracatinib (1 µM), EGFR inhibitor Gefitinib (10 µM), EGFR/Her2 inhibitor Lapatinib (10 µM), FAK inhibitor FAKi 14 (10 µM), BTK inhibitor Ibrutinib (1 µM) or Tec-family kinase inhibitor LFM-A13 (20 µM) for 1hr. Eluates were immuno-blotted with anti-p55g and anti phosphotyrosine (total) antibodies to assay endogenous phosphorylation of Y^258^XXM-PLEKHS1 with GFP and PLEKHS1 as input-controls (n=2, except results with PP2 which are n=3).

We used LNCaP cells to test some of our *in vivo* data and allow us to address some mechanistic questions. In Figure 3 we showed that when *mPlekhs1*-eGFP was co expressed with active c*SRC* in LNCaP cells it was phosphorylated on Y^258^ an associated with p55γ and p85 proteins; both of these events were inhibited by a PI3K inhibitor (PI-103) (Figure 5B), confirming our *in vivo* data. When mPLEKHS1-eGFP was expressed in the absence of heterologous Y527F-*cSRC* it was still possible to detect clear binding of p55γ (Figure 3B). This binding of p55γ was unaffected by overnight serum-starvation, suggesting it was not driven by serum-derived factors but reduced by a partially-selective inhibitor of SRC-family kinases (Figure 5C). We used it as a read-out of endogenous phosphorylation of Y^258^-PLEKHS1 to test the impact of a range of selective tyrosine kinase inhibitors (Figure 5D). This revealed that inhibitors that targeted SFKs, and not FAK-, BTK-, EGFR and EGFR/Her2 selective inhibitors, reduced binding of p55γ and p85s. These results demonstrate endogenous SFKs can phosphorylate Y^258^-mPLEKHS1 in LNCaP cells.

The expression and activity of SFKs in mouse prostate was substantially increased upon deletion of PTEN but insensitive to PI3K inhibitors (Figure S2C). These results strengthen the case that SFKs are responsible for phosphorylation of Y^258^-PLEKHS1 in PTEN-KO prostate. They also suggest it is unlikely that the sensitivity of Y^258^ PLEKHS1 phosphorylation to PI3K inhibitors is due to PI3K-driven activation of SFKs, but, rather, it is entirely explained by PIP_3_/PI(3,4)P_2_-driven, PH domain mediated relocalisation of PLEKHS1.

### PIP_3_ accumulation, AKT phosphorylation and growth in PTEN-null but not wild type prostate are dependent on PLEKHS1 in the prostate epithelium

To test whether PLEKHS1 was required for PIP_3_ pathway activation in PTEN-KO mouse prostate, a *Plekhs1*^-/-^ mouse model was derived using targeted ES cells (EUCOMM, *Plekhs1^tm2a(EUCOMM)Hmgu^*). *Plekhs1*^-/-^ mice were viable and without overt, prostate growth/morphology nor fecundity phenotypes (Figure S6A-D). The mice were interbred with *Pten^loxP/loxP^* and *PbCre4*^+^ strains to obtain relevant genotypes for experiments (immuno-blots confirmed PLEKHS1 and/or PTEN were lost from the prostates of these mice as expected, Figure S6E). Analysis of their prostates showed that PLEKHS1 was required for PIP_3_ accumulation, AKT phosphorylation, tissue dysplasia and growth in PTEN-KO, but not wild-type, tissue (Figure 6A-E).

**Figure 6.**
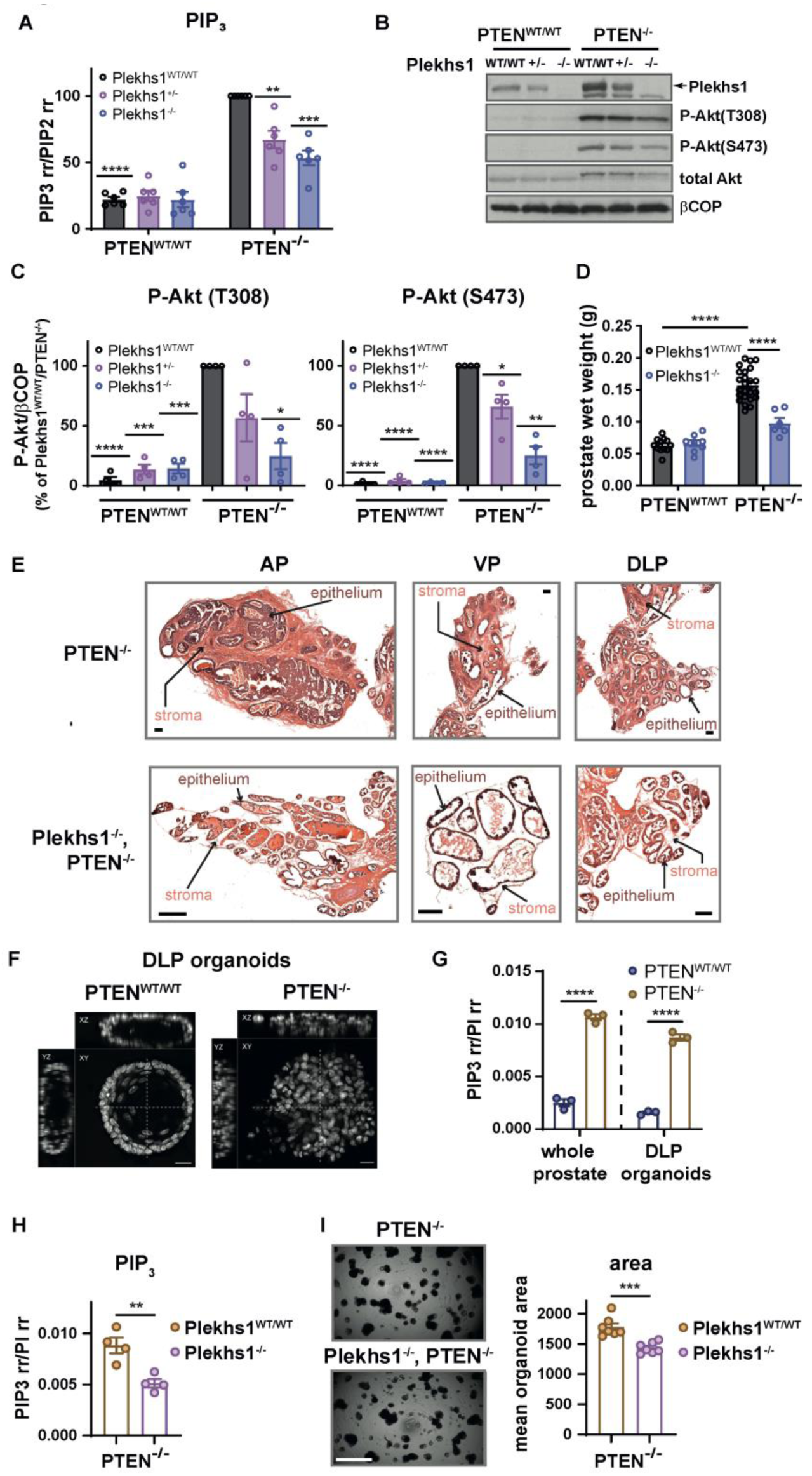
PIP_3_ accumulation, AKT phosphorylation and growth in PTEN-null but not wild-type prostate are dependent on PLEKHS1 in the prostate epithelium. Whole prostates were rapidly dissected from mice (12-15 weeks) with the genotypes indicated. **A**, Flash frozen material was extracted and total PIP_3_ was quantified, normalised to total PIP_2_ quantified in the same sample and presented as a % of the amount found in *Pten^-/-^ x Plekhs1^WT/WT^* samples. Data are means ± SEM of 6 biological replicates per genotype. For statistical analysis, multiple one-sample *t*-tests were performed, with Holm-Šídák’s correction Significant p value summaries (p≤0.05) are indicated. **B and C,** Frozen prostate was solubilised and P-T^308^ and P-S^473^-AKT were quantified relative to the amount of βCOP in the same samples by immuno-blotting. A representative immuno-blot is shown in (**B)**. **C,** The data are presented as a % of the *Pten^-/-^ x Plekhs1^WT/WT^* samples and are means ± SEM of n=4 biological replicates for each genotype. For statistical analysis, multiple one sample *t*-tests were performed, with Holm-Šídák’s correction. Significant p value summaries (p≤0.05) are indicated. **D,** Prostate wet weights from mice of the indicated genotypes. Data are means ± SEM (n=12, *Plekhs1^WT/WT^x Pten^WT/WT^*; n=8, *Plekhs1^-/-^x Pten^WT/WT^*; n=24, *Plekhs1^WT/WT^x Pten^-/-^* and n=6 *Plekhs1^-/-^x Pten^-/-^*, biological replicates). For statistical analysis, a 2way ANOVA was performed followed by Holm-Šídák’s multiple comparisons tests. Significant P value summaries (p≤0.05) are indicated. **e,** H&E-stained cryosections of prostates taken from the indicated lobes and genotypes. AP: anterior prostate; VP: ventral prostate; DLP: dorsolateral prostate. Representative images from n=3 biological replicates are shown. Scalebars: 200μm. **F, G, H and I,** DLP lobes from *Pten^WT/WT^* and *Pten^-/-^* mouse prostates (12-15 weeks) were used to derive organoids. **F**, DLP organoids were fixed, stained with DAPI to reveal nuclei and visualized by confocal microscopy. The images were constructed from Z-stacks of confocal sections and show a mid level section through the organoids. Scalebars: 20 μm. **G,** PIP_3_ measurements in whole prostate tissue and DLP organoids. Data are means ± SEM of 3 biological replicates per genotype and are expressed as the ratio of the abundance of total PIP_3_ to that of total PI in the same sample. For statistical analysis, a 2-way ANOVA was performed followed by Holm-Šídák’s multiple comparisons tests. Significant p value summaries (p≤0.05) are indicated. **H,** PIP_3_ measurements in *Pten^-/-^ x Plekhs1^WT/WT^* and *Plekhs1^-/-^ x Pten^-/-^* DLP organoids. Data are means ± SEM of 4 biological replicates per genotype and are expressed as the ratio of the abundance of total PIP_3_ to that of total PI in the same sample. **I,** Transmitted light images from *Pten^-/-^ x Plekhs1^+/+^* and *Plekhs1^-/-^ x Pten^-/-^* DLP organoids. A representative image is shown for each genotype. Data are mean ± SEM (n=7 biological replicates per genotype where each data point is an average area of 71-136 organoids). For statistical analysis of **H** and **I**, an unpaired, two-tailed T-test was performed. Significant p value summaries (p≤0.05) are indicated.

We addressed whether the PLEKHS1-dependency of PIP_3_-accumulation and growth in PTEN-KO mouse prostate was prostate epithelium autonomous. Organoids were derived from mouse prostates with relevant genotypes using published protocols (Drost et al., 2016). Organoids derived from PTEN-KO prostate grew more robustly and had substantially higher levels of PIP_3_ than PTEN-wild-type organoids and both phenotypes were dependent on PLEKHS1 expression (Figure 6F-I and S6F); indicating the phenotype of PLEKHS-KO mouse prostate is a direct result of loss of PLEKHS1 function in prostate epithelial cells.

Together, these results suggest that the high levels of PIP_3_ and PI(3,4)P_2_ in PTEN KO prostate epithelial cells depend upon cell-intrinsic PLEKHS1. In the context of our conclusion, that PIP_3_/PI(3,4)P_2_ binding at the PH domain of PLEKHS1 is required for phosphorylation of Y^258^ and hence activation of class I PI3Ks, this suggests that a self-sustaining loop may have been established.

### Levels of hPLEKHS1 mRNA and activating phosphorylation of Src-family kinases correlate with PI3K pathway activation in human prostate cancer

Previous studies have reported that PI3K/Akt/mTOR pathway activation can occur without proteogenomic associations with putative drivers or mutations in a large subset of prostate cancers (Cancer Genome Atlas Research, 2015; Zhang et al., 2017). Although *PTEN* deletion occurs commonly (∼17%) in primary prostate cancers, it correlates poorly with pathway activation in cohort analysis, likely due to confounding effects of alternate mechanisms of PI3K pathway deregulation (e.g., *SPOP* mutation) (Blattner et al., 2017; Cancer Genome Atlas Research, 2015). Lack of predictive biomarkers for pathway activation presents a major challenge for therapeutic PI3K targeting and patient stratification. In the context that h*PLEKHS1* is also expressed in both healthy and cancerous prostate, we investigated TCGA PRAD datasets for evidence of PLEKHS1-mediated PI3K activation. Although *PLEKHS1* mutations were rare (1%), *PLEKHS1* gene expression was significantly increased in primary prostate tumours compared to healthy tissue (Figure 7A and S7A). Changes in *PLEKHS1* mRNA expression were queried further as they correlated with protein levels in cancer types where proteomics data was available and, furthermore, we have shown simple over-expression of wild-type, but not Y^258^F-mutant, PLEKHS1 in LNCaP cells can increase PIP_3_ (Figure 3C). Grouped comparison of primary prostate cancers based on quartiles of *PLEKHS1* mRNA expression (Figure S7B) showed significant positive correlations with activation of key pathway proteins (Akt, 4E-BP1 and S6 phosphorylation) (Figure 7B and S7D, E). Furthermore, SRC-Y419-phosphorylation also correlated strongly with PI3K pathway activation in primary prostate cancers, in agreement with previous reports (Cancer Genome Atlas Research, 2015) (Figure 7C, S7C). Overall, these findings support the clinical relevance of a SRC-PLEKHS1-PI3K signalling axis in human prostate cancer cells.

**Figure 7.**
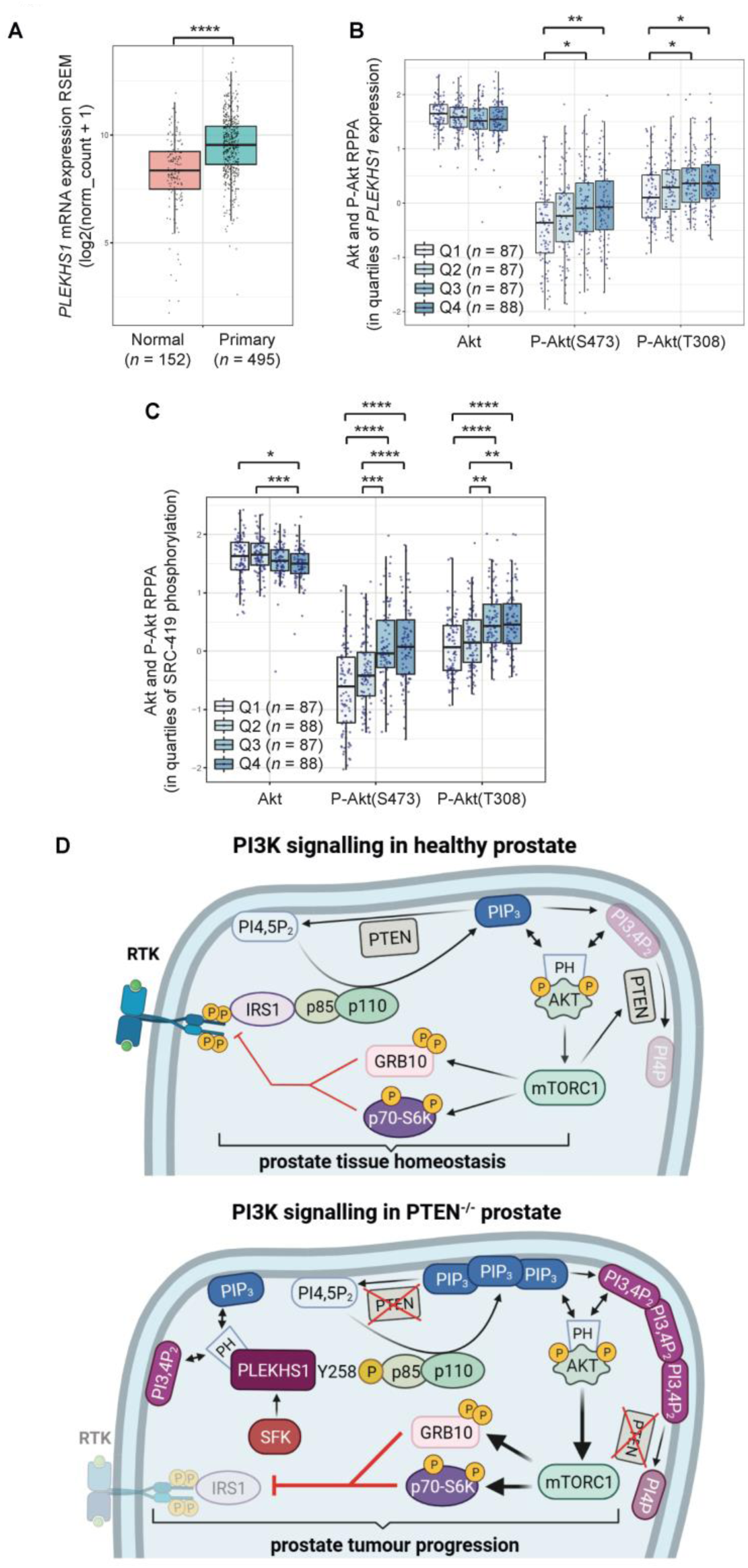
*PLEKHS1* expression and SRC-Y419-phosphorylation correlate with PI3K pathway activation and support a model for PI3K activation in PTEN-KO prostate. **A,** *PLEKHS1* mRNA expression (RSEM) in normal prostate and primary prostate cancer. Data from UCSC Xena database (TCGA PanCancer, TCGA Target, GTEx). Statistics: Welch’s t-test. **B,** Primary prostate cancer samples were grouped by quartiles of *PLEKHS1* mRNA expression (RSEM) and compared for Akt and p-Akt (S473, T308) protein levels (RPPA). Data from cBioPortal TCGA-PRAD. Statistics: Kruskal-Wallis with Dunn’s multiple comparison test. **C,** Primary prostate cancer samples were grouped by quartiles of p-SRC (Y419) protein levels (RPPA) and compared for AKT and P-AKT (S473, T308) protein levels (RPPA). Data from cBioPortal TCGA-PRAD. Statistics: Kruskal-Wallis with Dunn’s multiple comparison test. **D,** Model for PI3K signalling in healthy and *PTEN*^-/-^ prostate. In healthy prostate, PI3K signalling is mainly driven by IRS family members and fine-tuned by PTEN activity and two forms of feedback from mTORC1. Firstly, *via* p70s6K and Grb10, reduced expression of RTKs and IRS leading to reduced activation of PI3Ks. Secondly, *via* 4E-BP1, increased synthesis and accumulation of PTEN. In PTENs^-/-^ prostate, the PI3K signalling network is dramatically re-wired. PLEKHS1 becomes a major activator and target of PI3K signalling, sustaining pathway activity despite intense negative feedback. We propose that re-wiring of PI3K signalling underpins PTEN^-/-^ prostate tumour progression.

## Discussion

Our experiments allow us to suggest that in LNCaP cells a SFK is responsible for phosphorylation of Y^258^-PLEKHS1. Given that the SFKs most highly expressed in LNCaP cells are *YES >SRC > LYN >FYN* (Spans et al., 2014; Taberlay et al., 2016), presumably one or more of these members catalyses this reaction. Healthy mouse prostate tissue expresses significant amounts of five SFKs (*Src > Frk > Yes > Lyn > Fyn* mRNAs), the levels of three of which increase by about two-fold in PIN-stage, PTEN-KO tissue (*Src > Frk > Lyn* mRNAs) (Jurmeister et al., 2018). Our data show that there is a large increase in SFK expression and activity in PTEN-KO tissue. This increase in SFK activity may result from an increase in ROS, caused by loss of PTEN specifically (Li et al., 2013) and cancer progression more generally (Hayes et al., 2020), leading to their sulfenylation and activation (Heppner et al., 2018). These factors suggest SFKs are more active and play a significant role in coordinating a broad pattern of change in PTEN-KO mouse prostate and human prostate cancer and that part of their impact may be *via* phosphorylation of PLEKHS1.

A number of lines of evidence suggest that in the absence of PTEN the PI3K pathway could become relatively insensitive to the receptor mechanisms that are key determinants of its activity in healthy prostate: 1) the dramatic reductions in binding of p85s to IRS, EGFR and SHC1 (a receptor proximal adaptor) in PTEN-KO prostate; 2) the potential for PLEKHS1 to be subject to positive feedback from PI3Ks; 3) the lack of RTKs and their adaptors in the STRING-network of PLEKHS1 compared to GAB and IRS proteins; 4) the insensitivity of YXXM-phosphorylation of PLEKHS1 in LNCaP cells to serum-starvation; 5) work demonstrating that tumours with activating mutations in the PI3K pathway, including mouse prostate-specific loss of PTEN, are insensitive to dietary restriction (Kalaany and Sabatini, 2009) and, finally, 6) the possibility that SFKs could be activated by ROS in PTEN-KO prostate.

We propose a qualitative model (Figure 7D) that can rationalise our results. In healthy prostate the PI3K pathway is a key determinant of the growth of the tissue. Its activity is primarily driven by receptors capable of eliciting tyrosine phosphorylation of IRS proteins. Physiological pathway activity is constrained by a combination of PI(3,4)P_2_/ PIP_3_ phosphatases, particularly PTEN, and pathway feedback suppressing IRS-mediated activation of PI3Ks and augmenting PTEN expression. The outcome of these counter-poised factors establishes a homeostatic mechanism that shapes the output of the pathway. A key feature is sensitivity to extracellular growth factors such as insulin and IGFs. Upon loss of PTEN activity, levels of PI(3,4)P_2_/ PIP_3_ and the activity of downstream targets, such as AKT and MTOR, are increased. Pathway feedback is enhanced and suppresses expression and phosphorylation of IRS, limiting the increases in PI(3,4)P_2_/ PIP_3_. Some PI3K activators are unchanged by loss of PTEN. However, the increase in both PI(3,4)P_2_ and PIP_3_ induced by loss of PTEN is well-matched to the selectivity of the PH domain of PLEKHS1 and is likely to cause its localisation to membranes. SFKs are more active in PTEN-KO cells, in part as a result of ROS-induced sulfenylation and part *via* the loss of protein phosphatase activity of PTEN leading to de-repression of PTK6 and SRC (Alwanian et al., 2022, Wozniak et al., 2017). This leads to increased phosphorylation of PLEKHS1 on Y^258^ by SFKs and a dramatic further increase in activation of class I PI3Ks. As a result, and in the context of the suppression of IRS signalling and the insensitivity of PLEKHS1 to pathway feedback, PLEKHS1 becomes a major activator of the PI3K pathway. In this setting the PI3K pathway might become less dependent on growth factor drive to maintain its activity.

Our results also point to the intriguing possibility that PLEKHS1 signalling may be limited by association with 14:3:3 proteins. 14:3:3 proteins form dimeric scaffolds that can bind proteins phosphorylated on serine or threonine residues within preferred local sequences (Pennington et al., 2018). These interactions lead to many outcomes, including re-localisation, by revealing targeting motifs within the phosphorylated proteins, and separation of complexed proteins (Gardino et al., 2006). Three phosphoserine residues in PLEKHS1 are predicted to bind 14:3:3 proteins (Madeira et al., 2015) (ranked predicted sites: S^157^, S^256^ and S^180^) all of which are more heavily phosphorylated in PTEN-KO prostate (Figure S5A). S256 is also very close to the Y^258^ site that can bind to p85 and may therefore establish a competition between p85 and 14:3:3 binding, similar to the role of 14:3:3 proteins in the termination of signalling by GAB2 (Brummer et al., 2008). This suggests that PLEKHS1 is either complexed to 14:3:3 proteins or PI3Ks and that conclusion is supported by string analysis of the family of specific PLEKHS1 interactors we have defined (Figure 5A).

Another protein that was substantially more abundant in p85-pulldowns from PTEN KO mouse prostate is AFAP1L2 (XB130). It is a well-studied protein that is differentially expressed between tissues and is detected in mouse prostate. It harbours a Y^54^XXM motif, demonstrated to bind and activate PI3K signalling (Lodyga et al., 2009) and can be recruited to actin-rich regions of the cell periphery (Lodyga et al., 2010). The two PH domains of AFAP1L2 are unlikely to bind phosphoinositides, but AFAP1L2 interacts with SH3PXD2A (Moodley et al., 2015) (TKS5/FISH, expressed in mouse prostate) a PX-domain-containing adaptor that can bind PI(3,4)P2 and be recruited to podosomes (Abram et al., 2003). Numerous studies suggest AFAP1L2 can drive tumour progression (Zhang et al., 2016). Importantly, AFAP1L2 is a recognised interactor and substrate of SRC (Flynn et al., 1993). Taken together these observations suggest AFAP1L2 may also contribute to processes of pathway remodelling and tumour progression in PTEN-KO mouse prostate.

It is unclear why PLEKHS1 takes on such an important role in driving PI3K pathway activity in PTEN-null mouse prostate whilst the related GAB proteins apparently do not. GAB proteins have a similar architecture and regulatory properties (Rodrigues et al., 2000) to PLEKHS1 and can also be phosphorylated by SFKs (Kong et al., 2003). The observation that recovery of GABs with p85 and their expression are both broadly unchanged upon loss of PTEN may indicate they are not abundant in prostate luminal cells. The insensitivity of the interactions between GABs and p85s to PI3K inhibitors could suggest they are not sensitive to positive, PH-domain mediated feedback in this setting. Alternatively, although they can be substrates of SFKs, in this context and in contrast to PLEKHS1, they are not, because they fail to colocalise appropriately. Whatever the precise explanation, this re-enforces the need to study PI3K pathway activity in the *in vivo* context.

PLEKHS1 appears to be more widely expressed in human tissues than mouse and this may mean more cell types might gain advantage from the ability of PLEKHS1 to sustain chronic activation of the PIP_3_-network. This idea is supported by work that has linked PLEKHS1 to tumour progression in a range of tissues, including gastric, colorectal, hepatocellular, thyroid as well as prostate cancers. The majority of those studies positively-correlated *hPLEKHS1*expression or locus-demethylation with disease (Chen et al., 2021; Deng et al., 2019; Liu et al., 2020; Ruiz-Deya et al., 2021; Xing et al., 2020; Xiong et al., 2015; Zhang et al., 2020b) whilst another found a negative correlation (Abdel-Tawab et al., 2022). Consistent with the majority of those studies, heterologous over-expression of hPLEKHS1 has been reported to increase AKT phosphorylation and invasiveness in thyroid cancer cells (Xing et al., 2020). The apparent inconsistency in this data may be related to tissue-type and/or the apparently limited relevance of PLEKHS1 protein or mRNA levels compared to Y^258^-phosphorylation seen in our mouse work.

In terms of a potentially wider role for PLEKHS1, several studies have also reported a hot-spot of somatic, non-coding mutations at the apex of a palindromic-hairpin in the first intron of PLEKHS1 in cancer. These mutations are at their most frequent in bladder cancer where they occur in 40% of cases (Weinhold et al., 2014). Further studies have confirmed this observation and suggested it may result from coincident high expression of APOBEC activity in the cancer (Langenbucher et al., 2021; Rheinbay et al., 2020; Vacher et al., 2020; Wong et al., 2022). The presence of the mutations appears to correlate with reduced expression in some studies (Weinhold et al., 2014; Wong et al., 2022) but others, however, find no correlation in the same cancer type (Fredriksson et al., 2014). This has led different studies to classify the mutation as either a potential driver or passenger in tumour progression (Fredriksson et al., 2014; Wong et al., 2022). Differential expression of *Plekhs1*-mRNA between obese rats and obese diabetic rats, but not between control and diabetic rats, has been interpreted as indicating a role for PLEKHS1 in early progression towards type 2 diabetes (Kotoh et al., 2016). This study was part of the literature-based evidence used to argue the non-coding mutations in the *Plekhs1*-locus may interact with the IGF1 signalling axis to promote bladder cancer progression (Gill et al., 2021).

A clear implication of our work is that oncogenic, constitutive activation of the PI3K pathway can lead to a fundamental rewiring of its architecture. In this setting, remodelling is shaped by mechanisms that are insensitive to homeostasis and establish a higher, more cell-autonomous, set-point for pathway activity, thus priming the cell to take advantage of further oncogenic mutations. This points to the potential for identifying new vulnerabilities in the PI3K network in cancer cells.

## Supporting information

Chessa et al Supplementary Information

## Acknowledgements

The work was supported by grants from the BBSRC (BB/P013384/1 and BB/P020240/1), the MRC (MR/R000409/1), an ITN PhD studentship (to PJ), a CRUK Cambridge Center PhD (AA) and a TAG grant from the KEC programme at the Babraham Institute (BI).

We would like to thank; Professor Vincent Gnanapragasam (Translational Prostate Cancer Group, CRUK Cambridge Cancer Centre, UK), Clive D’Santos (Proteomics, CRUK, Cambridge Institute), Antonio Ramos-Montoya and Ajoeb Baridi (CRUK, Cambridge Cancer Centre, at that time) for helpful discussions; Estere Seinkmane for culturing mouse prostate organoids; the staff in BSU at BI for mouse work; Judith Webster for preparation of samples for mass spectrometry analysis; Keith Davidson (BI, Signalling) for technical assistance; Attila Bebes, Aleksandra Lazowska-Addy and Rachael Walker (BI, Flow Facility) for help with cell sorting experiments; Elena Ivanova (BI, Epigenetics) for help cryo-sectioning; Felix Krueger (BI, Bioinformatics, at that time) for help analysing mRNA-seq data and Chiranjeevi Sandi (AZ, Cambridge, at that time) for sharing unpublished mRNA-seq data.

## Author contributions

T.C., contributed to conception of the project, performed and analysed experiments, prepared figures and wrote the manuscript; P.J., Created GM-PLEKHS1 strains, performed experiments and analysed their data; S.S., performed experiments, analysed data, prepared figures and text for the manuscript; A.A., performed experiments, analysed data, prepared figures and text for the manuscript; K.A., performed lipid analysis; D.B., optimised FACS-sort experiments with cells from mouse prostate; A.K., performed lipid analysis; B.A.S., prepared an anti-Avi monoclonal antibody and prepared organoids; S.F., analysed histochemical slides; D.O., proteomic workflow and data analysis; D.S., generation of all GM mouse models; A.S-P., provided statistical expertise and generated the heat map, M.W., provided structural insight into PLEKHS1 protein and domain expression; S.W., performed confocal analysis of organoids; H.O., created image analysis pipelines to quantify organoid area; S.C., contributed to grant-writing, planning and writing of the manuscript; P.T.H. and L.R.S., conceived the project, wrote grants, planned experiments, interpreted data and wrote the manuscript.

## STAR Methods

### RESOURCE AVAILABILITY

#### Materials availability

Plasmids and mouse models generated in this study will be made available upon request.

#### Lead contact

Correspondence and requests for materials should be addressed to and will be fulfilled by; Tamara Chessa, or Len Stephens, the Babraham Institute, Cambridge, CB22 3AT, UK.

#### Data availability

The majority of data generated and analysed by this study are included in the published article and its supplementary information files. TMT-LC-MS/MS data will be made available on ProteomeXchange.

### EXPERIMENTAL MODEL AND SUBJECT DETAILS

#### Experimental animals

*Pik3r1^Avi/Avi^*, *Pik3r2^Avi/Avi^* and *mBirA^+/+^* mice(Tsolakos et al., 2018) were bred with *PbCre4^+^* mice(Wu et al., 2001) and *Pten^loxP/loxP^* mice(Trotman et al., 2003) to generate *p85^WT/WT^* x *Pten^WT/WT^* (*PbCre4^-^* x *mBirA^+/-^* x *Pten^loxP/loxP^*), *p85α^Avi/Avi^ x Pten^WT/WT^* (*PbCre4^-^* x *Pik3r1^Avi/Avi^* x *mBirA^+/-^* x *Pten^loxP/loxP^*), *p85β^Avi/Avi^ x Pten^WT/WT^* (*PbCre4^-^* x *Pik3r2^Avi/Avi^* x *mBirA^+/-^* x *Pten^loxP/loxP^*), *p85^WT/WT^ x Pten^-/-^* (*PbCre4^+^* x *mBirA^+/-^* x *Pten^loxP/loxP^*), *p85α^Avi/Avi^ x Pten^-/-^* (*PbCre4^+^* x *Pik3r1^Avi/Avi^* x *mBirA^+/-^* x *Pten^loxP/loxP^*), *p85β^Avi/Avi^ x Pten^-/-^* (*PbCre4^+^* x *Pik3r2^Avi/Avi^* x *mBirA^+/-^* x *Pten^loxP/loxP^*).

Resulting mice were backcrossed to the C57BL/6J strain for at least 4 generations. For breeding, only *PbCre4^+^* males were used.

*Plekhs1^Avi/Avi^* mice were generated in BI facilities using CRISPR/Cas9 technology. gRNAs were designed with the Guide RNA Design Tool (www.benchling.com). Zygotes used to generate *Plekhs1^Avi/Avi^*mice were isolated from *Pten^loxP/loxP^*, *BirA^+/+^*females and then transferred to a WT C57BL/6J foster mother. Mice were bred with *PbCre4^+^*, *Pten^loxP/loxP^* mice to generate PLEKHS1-Avi x *Pten^WT/WT^* (*PbCre4^-^* x *Plekhs1^Avi/Avi^* x *mBirA^+/-^* x *Pten^loxP/loxP^*), PLEKHS1-Avi x *Pten^-/-^* (*PbCre4^+^* x *Plekhs1^Avi/Avi^*x *mBirA^+/-^* x *Pten^loxP/loxP^*) mice and their appropriate no-Avi controls.

*Plekhs1^-/-^* mice were generated using two genetically engineered embryonic stem cells clones purchased from Eucomm (Clone IDs HEPDO817_1_F11, HEPDo817_1_D12; Allele *Plekhs1^tm2a(EUCOMM)Hmgu^*) injected to a zygote isolated from *Pten^loxP/loxP^* females and then transferred to a WT C57BL/6J foster mother. Mice were bred with *Pten^loxP/loxP^* mice and *PbCre4^+^*, *Pten^loxP/loxP^*mice to generate *Plekhs1^WT/WT^* x *Pten^WT/WT^*, *Plekhs1^+/-^* x *PTEN^WT/WT^*, *Plekhs1^-/-^*x *Pten^WT/WT^*, *Plekhs1^WT/WT^* x *Pten^-/-^*, *Plekhs1^+/-^* x *PTEN^-/-^* and *Plekhs1^-/-^*x *Pten^-/-^* mice.

#### Experimental procedures

To determine effects of the pan-PI3K inhibitor GDC-0941 on levels of PIP_3_, p85-PLEKHS1 interactions, PLEKHS1 phosphorylation and growth factor signalling *in vivo* we treated *p85α^Avi/Avi^ x Pten^WT/WT^*, *p85α^Avi/Avi^ x Pten^-/-^,* PLEKHS1-Avi x *Pten^WT/WT^* (*PbCre4^-^*x *Plekhs1^Avi/Avi^* x *mBirA^+/-^* x *Pten^loxP/loxP^*) and PLEKHS1-Avi x *Pten^-/-^* mice with 800 mg/kg GDC-0941 formulated in HPMC/Tween [0.5 % hydroxypropyl methocellulose (Methocel (Colorcon))/0.2 % Polysorbate80] or vehicle p.o. once daily, for 2 days. Animals were sacrificed between 2 and 5 hours after administration of the last dose, at which plasma concentration of GDC-0941 was expected to be between 5 and 10 μM(Yang et al., 2016).

#### Ethical statement

All animal experiments at The Babraham Institute were reviewed and approved by The Animal Welfare and Ethics Review Body and performed under Home Office Project license PPL 70/8100.

#### Housing and husbandry

All animals used in this study were housed in the Biological Support Unit at the Babraham Institute and kept under specific pathogen–free conditions. Animals housed in the Biological Support Unit at the Babraham Institute were kept under specific pathogen–free conditions.

#### Animal care and monitoring

The animals were kept under SPF conditions and the animal facilities where the mice were kept were regularly checked for standard pathogens. The mice were looked after by professional caretakers. Every animal was checked daily. Health reports can be provided upon request.

Food and water were provided ad libitum. The light cycle ran from 6 am to 6 pm.

#### Study design, Sample size and Randomisation

Prostates from several age-matched male mice of identical genotype, randomly allocated to different experimental groups, were analysed, as detailed in the legends to figures. Multiple independent experiments were carried out using several biological replicates, as detailed in the legends to figures.

#### Cell Lines

##### LNCAP

LNCAP cells were obtained from the AstraZeneca cell bank and had been previously authenticated using DNA fingerprinting short tandem repeat assays. Cells were cultured in RPMI-1640 supplemented with 10% FBS and 1% w/v penicillin/streptomycin, for a maximum of 25 passages post-thaw.

##### Organoid culture

Prostate cell isolation and subsequent organoid culture was performed according to methodology described by Karthaus *et al*.(Karthaus et al., 2014). For organoid culture maintenance and imaging, cells derived from dorsolateral prostate (DLP) lobes were cultured by overlaying 10000 cells, in 400 μl full organoid growth medium (DMEM/F12 or ADMEM/F12 supplemented with 10 mM HEPES pH 7.4, 2 mM Glutamax and 1x Penicillin Streptomycin, as well as additional components B27 minus vitamin A, N-acetyl-L-cysteine (1.25 mM), murine EGF (50 ng/ml), recombinant murine Noggin (100 ng/ml), recombinant human R-Spondin-1 (500 ng/ml), Y-27632 dihydrochloride (10 μM), 5α-Dihydrotestosterone (DHT) solution (1 nM) and A 83-01 (200 nM), onto 95 μl Corning Matrigel Growth Factor Reduced (GFR) Basement Membrane Matrix, in a Nunc LabTek II chamber slide system (4 well; glass).

For PIP_3_ measurements, 5000 cells in 200 μl full organoid growth medium were cultured in 96-well plates containing 47.5 μl growth factor-reduced Matrigel for 4 days. Organoids were cultured for a maximum of 8 passages in complete ADMEM/F12 medium (Figure 5g) or for 1 passage in complete DMEM/F12 medium (Figure 5h).

For transmitted light imaging, 5000 cells in 200 μl complete ADMEM/F12 medium were cultured in 96-well plates for 6 days. For confocal imaging, organoids were cultured in a Nunc LabTek II chamber slide. Organoids were cultured in complete ADMEM/F12 medium for a total of 3 passages (Figure 5f), or imaged at passage 0 (Figure 5i).

### METHOD DETAILS

#### Prostate isolation and streptavidin-mediated pulldown of avi-tagged proteins

Mice were sacrificed using Schedule 1 methods and prostates rapidly dissected, rinsed in PBS, flash-frozen in N_2_(l) and stored at -80℃ until use. Tissues were pulverised under continuous flow of N_2_(l). For streptavidin-mediated pulldown, typically, 750ul Triton/CHAPS lysis buffer (1 % Triton X-100, 0.4 % CHAPS, 20 mM Tris-HCl pH 7.4, 150 mM NaCl, 1 mM EDTA, 1 mM EGTA, 2.5 mM Na_4_P_2_O_7_, 25 mM NaF, 5 mM Na_3_VO_4_, 5 mM β-glycerophosphate, 10 μg/ml each of leupeptin, aprotinin, α−pain and pepstatin A, 1 mM PMSF) was added to 50 mg of pulverised tissue to yield an estimated protein concentration of 4 mg/ml. For *Pten^-/-^* prostate, depending on variations in fluid content of the tissue, a correction factor had to be applied. Lysates were cleared by ultracentrifugation (40000 rpm, 5 minutes, Beckman Optima Max centrifuge, MLA-130 rotor). An estimated total of 2 mg protein was used per pulldown.

For PLEKHS1-Avi pulldowns followed by immunoblotting with anti-phospho-Y^258^ PLEKHS1 antibody, prostates were lysed in a modified RIPA buffer (50 mM Tris-HCl pH 7.4, 1% NP-40, 0.5 % sodium deoxycholate, 0.1 % SDS, 150 mM NaCl, 1 mM EDTA, 1 mM NaF, 2.5 mM Na_4_P_2_O_7_, 5 mM Na_3_VO_4_, 5 mM β-glycerophosphate, 10 μg/ml each of leupeptin, aprotinin, α−pain and pepstatin A, 1 mM PMSF). An estimated total of 6 mg protein was used per pulldown.

Broadly, streptavidin-mediated pulldown of Avi-tagged proteins was performed as described in Tsolakos et al.(Tsolakos et al., 2018) with the following modifications: total time of pulldown was 20 minutes and beads were washed 4 times post-pulldown prior to elution in 1.5 x SDS sample buffer. For pulldowns followed by immuno-blotting with anti-phospho Y^258^-PLEKHS1 antibody, total time of pulldown was 1 hour and washes were performed with increasing level of stringency: 3 consecutive washes with each of the following: modified RIPA with 0.5 % SDS, 0.5 M NaCl or no NaCl.

Biotinylated proteins were eluted from beads by incubation with 1.5 x (2 mg protein) or 2 x (6 mg protein) reducing SDS sample buffer (95 °C, 10 minutes, with 1x vortex after 5 minutes)

#### TMT-LC-MS/MS

TMT-6-plex (identification of p85α and -β^Avi/Avi^ interactome), TMT-10-plex and TMT-11 plex Isobaric Label Reagents (analysis of the effect of in vivo PI3K inhibition on the p85α- avi interactome; identification of *Plekhs1^Avi/Avi^* interactome) were from ThermoFisher Scientific. TMT-LC-MS/MS, protein identification and quantification was performed essentially according to Luff et al., 2021(Luff et al., 2021).

#### p85^Avi/Avi^ interactome analysis

Quantitative MS data (scaled abundances calculated in Proteome Discoverer) was analysed in RStudio. Abundance values for high-confidence proteins that were identified in at least 3 biological replicates per condition were transformed (sqrt) to meet the assumption of normality for parametric testing. To distinguish specific p85α and/or p85β-interactors from non-specific background, multiple two-sided unpaired *t*-tests (Holm correction) were performed. A stringent significance threshold (p<0.01) was applied to identify a total of 24 proteins with increased detection above *p85^WT/WT^*controls, and FC of 2 and above, were identified.

#### Plekhs1^Avi/Avi^ interactome analysis

Quantitative MS data (grouped abundances, where each protein is expressed as a proportion of the total signal measured for that protein across all replicates and genotypes, average set to 400, and calculated in Proteome Discoverer) were analysed in Excel. To distinguish specific PLEKHS1-interactors from non-specific background, ANOVA-based statistical tests were applied to calculate adjusted P value, significant interactors are defined by adjusted P value ≤ 0.05 and a fold change >2, for *Plekhs1^Avi/Avi^ x Pten^WT/WT^ : Plekhs1^WT/WT^ x Pten^WT/WT^* or *Plekhs1^Avi/Avi^ x Pten^-/-^ : Plekhs1^WT/WT^ x Pten^-/-^.* The data for all genotypes is based on n=3 biological replicates, except *Plekhs1^WT/WT^ x Pten^WT/WT^*, which is based on 2 biological replicates (as a result of the restrictions placed by 11-plex TMT labelling available at the time).

For phosphorylation analysis, phospho S/T/Y were included as additional variable modifications for the database search with Sequest HT, within Proteome Discoverer. The mass spectra of the PLEKHS1 phosphopeptides reported by Sequest were manually interpreted to determine phosphorylation sites.

#### Immunoprecipitation

*Pten^WT/WT^* and *Pten^-/-^* whole prostate lysates were prepared in Triton/CHAPS lysis buffer at an estimated protein concentration of 4 mg/ml as described in ‘Prostate isolation and streptavidin-mediated pulldown of avi-tagged proteins’. For p85α IP and control (βCOP), 10 μg antibody was pre-coupled to 50 μl Dynabeads Protein G according to manufacturer’s conditions and washed once in Triton/CHAPS pre-IP wash buffer (1% Triton X-100, 0.4 % CHAPS, 20 mM Tris-HCl pH 7.4, 150 mM NaCl, 1 mM EDTA, 1 mM EGTA, 2.5 mM Na_4_P_2_O_7_, 25 mM NaF). For phosphor-tyrosine IP, 4G10 Platinum Anti-Phospho-tyrosine Agarose Conjugate antibody was used (80 μl). Beads were washed 3x in PBS, followed by 3x in pre-IP wash buffer. All lysis and wash steps were performed using ice-cold buffers, on ice.

Lysates were incubated with beads for 90 minutes, end-on-end (19 rpm) at 4 °C. Beads were washed 4x in Triton/CHAPS lysis buffer and bound proteins were eluted in 40 μl 1.5x SDS sample buffer (Dynabeads) or 50 μl 2x SDS sample buffer (Agarose beads), at 95 °C, for 10 minutes, with 1x vortex after 5 minutes.

#### GFP-trap® pulldown assay

LNCAP cells cultured in 10 cm dishes were transfected 24 hr after seeding using Lipofectamine 3000 according to manufacturer’s protocol. Cells were harvested 48 hr after transfection in Triton/CHAPS lysis buffer (750 μl/dish). For inhibitor treatments, media was replaced and cells pre-treated for 1 hr before harvesting. For serum starvation, media was replaced with complete media or starvation media (RPMI-1640 with Pen-Strep, phenol-red free) 16 hrs before harvesting. Crude lysates were vortexed for 10 sec and incubated on ice for 10 min with brief vortex at 5 min. Lysates were cleared by centrifugation (>13,000 rpm, 10 min, 4 °C, bench centrifuge). ChromoTek GFP-trap® beads (30 μl/sample) were pre washed with lysis buffer (3 x 500 μl). Pulldown was performed by incubating cleared lysates (∼1.5 mg) with beads for 1 hr with rotation (19 rpm) in cold room. Aliquots of lysates pre and post-pulldown were reserved for western blotting and BCA assay (pre). Beads were washed with lysis buffer (4 x 1 mL) and bound proteins eluted by boiling for 10 min at 95 °C in 2x sample buffer (30 μl/sample). Samples were vortexed briefly before and during (5 min) elution. Pre-/Post-pulldown samples were boiled for 5 min at 95 °C in 4x sample buffer. Samples were snap frozen and kept in -80 °C for western blots.

#### Generation of custom-made antibodies

Monoclonal antibodies against the avi-tag were developed by Babraham Bioscience Technologies (BBT)/BRC technology development lab using established methodology(Kohler and Milstein, 1975), where mice were immunised with avi-tag peptide GLNDIFEAQKIEWHE.

Polyclonal antibodies against mouse P-Plekhs1 Y258 were generated by Cambridge Research Biochemicals, where rabbits were immunised with antigen peptide C]-AESN-[pY]-VS-Nle RS-amide. Harvest bleeds were purified against depletion peptide [C]-AESNYVS-Nle-RS amide, then the unbound material was purified against antigen.

Polyclonal antibodies against human PLEKHS1 were generated by Cambridge Research Biochemicals. Rabbits were immunised with antigen peptide sequence [C] APKRSPAIKKSQQKGARE-acid and purified by affinity chromatography on Thiopropyl Sepharose coupled with the antigen.

#### Western blot

Prostate lysates were prepared as described in section ‘Prostate isolation and streptavidin mediated pulldown of avi-tagged proteins’. LNCaP lysates and GFP-trap eluates were prepared as described in section ‘GFP-trap pulldown assay’. Lysates were heated at 95 ℃ for 5 minutes in 4x SDS sample buffer, amounts of protein as indicated in figure legends were resolved by SDS-PAGE, and transferred to PVDF membranes. Membranes were immunoblotted with the indicated primary antibodies at 4 °C overnight, followed by 1-2 h at room temperature. They were then washed in TBS (40 mM Tris/HCl, pH 8.0, 22 °C; 0.14 M, NaCl) containing 0.1 % v/v Tween 20 (TBST) and incubated with HRP-conjugated secondary antibodies. Membranes were washed in TBST, signals were detected by ECL and quantified using Licor Image Studio Lite v.5.2.

#### Imaging

Prostates, consisting of anterior, ventral, and dorsolateral lobes (one pair of each lobe), were dissected intact as described previously(Malek et al., 2017). 20 μm cryosections were prepared on charged glass slides using a Leica CM1850 cryostat. H&E staining was performed using Mayer’s hematoxylin solution and Eosin Y solution, following a standard protocol. Images were acquired using a Zeiss Axio Imager Z2 microscope (EC Plan-Neofluar 10x objective; AxioCam MR Rev3), and automatically stitched with AxioVision software, with shading correction enabled. 12 μm *Pten^WT/WT^* and *Pten^-/-^* prostate cryosections prepared on charged glass slides were rehydrated in PBS at room temperature for 10 minutes.

Cryosections were washed 1x and incubated for 1 hour in block/perm buffer (10 % horse serum, 1 % fatty acid and endotoxin-free BSA and 0.3 % Triton X-100 in PBS). Sections were then labelled with 10 μM Hoechst 33342 (prepared as a 1 mM solution in 200 ml tissue culture-grade dH_2_O and 2 ml 95 % ethanol) and CK8 antibody (1:500) in block/perm buffer, for 16 hours at 4 °C. Sections were washed 3x in PBS, followed by incubation with Goat anti-Rabbit IgG Antibody, Alexa Fluor 488. Sections were washed 3x in PBS, mounted in VECTASHIELD (#H-1000) and visualised on an EVOS M5000 cell imaging system, using the 20 x objective.

For *Pten^WT/WT^* and *Pten^-/-^* DLP organoid imaging, organoids were washed in 300 μl PBS and fixed in 200 μl 4 % paraformaldehyde in PBS for 20 minutes at room temperature. Organoids were washed in PBS and permeabilised with 200 μl per well PBS/1 % Triton X-100 (60 minutes at room temperature). Organoids were then blocked in 200 μl PBS/2 % BSA/1 % Triton X-100 per well (60 minutes at room temperature). Organoids were mounted in VECTASHIELD containing DAPI (diluted 1:3 in PBS) and imaged using a Nikon A1-R confocal microscope (Nikon Ti2 body) with a 20x 0.75 NA objective and operated using Nikon Elements software.

For quantification of DLP organoid area, transmitted light images were acquired on the EVOS M5000 microscope, using the 5x objective. Images were imported in Napari, and masks for all organoids generated using the Cellpose plugin (cellpose.org(Stringer et al., 2021)). Masks were corrected manually where necessary.

#### <u>Measurement of PIP</u>_3_

Pulverised prostate tissue (5 mg) was resuspended in initial organic solvent mix (chloroform:methanol 1:2) containing H_2_0 (organic mix:H_2_0 in 725:170 ratio). Following this step, 200 μl of the resuspended tissue was mixed with 720 μl of initial organic solvent mix:H_2_0 with added standards (d6-S/A-PIP_3_ (10 ng) and d6-SA-PI(4,5)P_2_ (100 ng)). Lipid extraction and quantification of levels of PIP_3_ was performed according to previously described methodology(Rynkiewicz et al., 2020).

LNCaP cells cultured in 6-well plates were transfected 24 hr after seeding using Lipofectamine 3000 according to manufacturer’s protocol. Cells were harvested 48 hr after transfection in 1 M HCl (750 μl/well). Lipid extraction and quantification of levels of PI(3,4,5)P3 was performed according to previously described methodology(Clark et al., 2011).

Organoids cultured in 96-well plates were harvested in 1 M HCl (250 μl/well). Lipid extraction and quantification of levels of PIP_3_ was performed according to previously described methodology(Clark et al., 2011).

#### Cloning of Plekhs1 from mouse prostate and generation of bacterial and mammalian expression constructs

Bacterial expression vector pRSET-2x(10His)-MBP-TEV site-mEGFP (hereafter referred to as pRSET-mEGFP) was generated using PCR and multiple insertions between the *Xba*I and *Nde*I sites of pRSET A vector (Thermo Fisher Scientific Waltham, MA, USA). Also, a protein-of-interest (POI)-linker-EGFP sequence was inserted between the *Nde*I (POI start), *Bam*HI (linker) and *Hind*III (EGFP stop) sites using multiple PCR and insertions.

RNA was extracted from whole mouse prostate of a C57BL/6J strain using TRIzol™ Reagent, according to manufacturer’s instructions. First-Strand cDNA synthesis was performed using SuperScript™ II Reverse Transcriptase in combination with Random Hexamer Primers. 20 μl of resulting cDNA was then used to amplify the isolated PH domain (iPH) of m*Plekhs1* (nucleotide sequence corresponding to amino acids 10-135), or full length m*Plekhs1* (nucleotide sequence corresponding to amino acids 10-459), followed by cloning into pRSET-mEGFP. For cloning of m*Plekhs1* iPH, primers containing engineered restriction sites *Nde*I and *Bam*HI were used. For insertion of full length mPlekhs1, the *Bam*HI linker site was modified by PCR to *Kpn*I.

The cloned sequence differs from the canonical m*Plekhs1* sequence (NCBI mouse RefSeq isoform 1), as follows: Serine 77: missing; Amino acids 323-336: missing. Hence, YxxM is Y257 in the cloned sequence (Fig 3a,b; 4c,d; Extended data 6b,c) and Y258 in the canonical sequence. The peptide coverage from PLEKHS1-Avi pulldowns from *Plekhs1^Avi/Avi^ x Pten^WT/WT^* and *Plekhs1^Avi/Avi^ x Pten^-/-^* prostates demonstrates that the canonical sequence is the most abundant in mouse prostate. For simplicity, we refer to YxxM in the main text as Y258. The 2x(10 His)MBP-mEGFP (pRSET-mEGFP) empty vector was generated by excision of POI (full length m*Plekhs1*)-linker-EGFP. mEGFP was amplified using primers with engineered restriction sites *Nde*I and *Hind*III, followed by cloning into pRSET-2x(10His) MBP-TEV site.

For mammalian expression, mouse *Plekhs1* cDNA was amplified from above mentioned pRSET-mEGFP constructs, using primers containing engineered restriction sites *Not*I (forward primer; this primer also carries KOZAK sequence, as well as sequence encoding amino acids 1-9 that are absent in pRSET-mEGFP constructs) and *Xba*I (reverse primer), and cloned into pCMV3-EE(Stephens et al., 2001). pCMV3-EE-mEGFP was generated by amplification of mEGFP from pRSET-mEGFP, using primers containing engineered restriction sites *Not*I and *Xba*I, followed by insertion into pCMV3-EE.

Site-directed mutagenesis to generate *Plekhs1*-Y257F constructs was performed with the Quikchange Lightning Site-directed Mutagenesis kit (Agilent), according to manufacturer’s instructions.

#### Recombinant protein expression and purification

Mouse PLEKHS1 (10-459) and its isolated PH domain (10-134) were expressed and purified as 2x(10His)-MBP-TEV site-(PLEKHS1 construct)-mEGFP) fusion proteins, along with 2x(10His)-MBP-TEV site-mEGFP alone, in BL21-CodonPlus (DE3)-RIPL Competent Cells. Cells were lysed by probe sonication on ice (0.1 M NaCl, 20 mM TRIS/HCl, 50 mM Na/HPO4, pH 7.5 with anti-proteases lacking EDTA (HALT^TM^) with the addition of TX100 to 1 % following sonication) and the lysates were cleared by centrifugation at 30,000 x g for 20 min. The proteins were purified *via* Co^2+^-affinity resin (His Pur^TM^ Cobalt resin), eluted by the addition of 0.25 M imidazole and buffer-exchanged *via* PD10 columns into 0.2 M NaCl, 1 mM EDTA, 20 mM HEPES/NaOH pH 7.4, at 25 ^0^C. For the liposome-based assays, the 2x(10His) MBP-tag was cleaved by incubation with 6xHis-tagged-Tobacco Etch Virus (TEV)-protease and repurified *via* Co^2+^ affinity resin. For the PIP strips assays, the proteins were further purified (from the Co^2+-^column eluates in 0.25 M imidazole) through their MBP tag (MBP TRAP, equilibrated in 0.2 M NaCl, 1 mM EGTA, 25 mM Na/HPO_4_ buffer pH 7.5 and eluted by the addition of 20 mM maltose). All proteins were concentrated (Amicon, 10 kD cut-off) and stored in 50 % (w/v) glycerol at -20 °C. Their purity and protein concentration (relative to BSA, in the range 0.5-2.0 mg/ml) were assessed by SDS PAGE and Coomassie staining.

#### Lipid binding assays

To test the lipid binding properties of PLEKSH1 and its iPH domain, protein-lipid overlay experiments were performed using PIP Strips^TM^ and PIP Array. Strips were blocked for 1 hr at RT in 3 % (w/v) BSA in TBST. PIP strips and arrays were then incubated for 1 hr in the blocking solution containing; 2.5 µg total of purified 2x(10His)-MBP-TEV site-PLEKSH1 mEGFP or 2x(10His)-MBP-TEV site-PLEKSH1-iPH-mEGFP or the control 2x(His10)-MBP TEV site-mEGFP proteins. After washing, the membranes were incubated for 1 hr with an anti GFP antibody in blocking buffer followed by washing with blocking buffer and then incubation with HRP goat anti-mouse antibody diluted in blocking buffer. The immunoreactive spots were detected using enhanced chemiluminescence.

For liposome-based sedimentation assays, liposomes were made by sonication of vacuum dried lipids, in quantities appropriate to give final concentrations in the assay of 150 mM phosphatidylserine, 200 mM phosphatidylcholine, 20 mM phosphatidylethanolamine, 10 mM sphingomyelin with various concentrations of PI(4,5)P_2_, PI(3,4)P_2_ or PI(3,4,5)P_3_ (0.1-4 mM or 0.026-1 mol%), into 0.2 M sucrose containing 20 mM KCl, 20 mM Hepes/NaOH, pH 7.4 at 25°C (the primary sonicated suspensions were diluted 10x into the final assay so that the final extra-liposomal concentration of sucrose was 20 mM). The liposomes were incubated with 120 ng of recombinant protein in assay buffer (total final volume of 150 ml, final concentrations in the assay of; 1 mg/ml BSA, 0.12 M NaCl, 1 mM EGTA, 0.2 mM CaCl_2_, 1.5 mM MgCl_2_, 18 mM Hepes/NaOH 7.4, 25 ^0^C; with approximately 100 nM free Ca^2+^) for 3 min at 30 °C and then pelleted by centrifugation at 40,000 × g for 5 min (Optima, benchtop ultracentrifuge, Beckman) Aliquots of the assays were taken before and after centrifugation, resolved by SDS PAGE and immuno-blotted (1^0^ antibody anti-GFP, 2^0^ anti-mouse-HRP) to quantify the proportion of the proteins that were sedimented with the sucrose-loaded liposomes.

#### Flow cytometry

Up to 4 PTEN^WT/WT^ or 2 PTEN^-/-^ prostates were digested in 4 ml Collagenase Type II, at a concentration of 5 mg/ml in ADMEM/F12 mix (ADMEM/F12 supplemented with 10 mM HEPES pH 7.4, 2 mM Glutamax and 1x Penicillin-Streptomycin) for 2 hours at 37 ℃, followed by further digestion in 2 ml TrypLE™ Express Enzyme with the addition of 10 μM Y-27632 for 15 minutes at 37 ℃. The resulting cell suspension was centrifuged (5 minutes, 350 RCF), resuspended in 1 ml PBS/0.5 % BSA, and passed through a 100 μm cell strainer to eliminate clumps of cells. To identify live, nucleated cells the single cell suspension was incubated with 10 μM Hoechst 33342 for 30 minutes at 37 ℃. Cells were incubated with CD326/EpCAM FITC (1:300) and CD49f-PE (1:200) at 4 ℃, for 45 minutes. Cells were centrifuged, washed in PBS/0.5 % BSA and resuspended in 330 μl sorting buffer (PBS/0.5 % BSA, 10 mM HEPES, 2 mM EDTA). 30 μl was reserved as an unsorted sample. 50 μl of TO-PRO3 was added to the remainder and cells were sorted into basal epithelial cells (defined as EpCAM+ CD49f-high), luminal epithelial cells (EpCAM+ CD49f-low) and remainder (EpCAM-) on a BD Influx Cell Sorter. The FITC fluorochrome was detected with the 488 nm laser and 530/30 bandpass filter, and the PE fluorochrome was detected with the 561 nm laser and 585/29 bandpass filter. During sorting, cells were kept at 4 ℃. Cells were collected in ice-cold PBS, a sample reserved for counting, centrifuged, and immediately flash-frozen in N_2_(l).

Cells were sorted on the BD Influx cell sorter using a 100 μm nozzle at 30 psi sheath pressure. 1 drop pure sort mode was used for collection into 1.5 ml tubes.

#### Bioinformatics analysis

*PLEKHS1* mRNA expression RNA-seq data (RNA Seq V2 RSEM) in normal prostate and primary prostate cancers was downloaded from UCSC Xena database (TCGA PanCancer, TCGA Target, GTEx).

Grouped comparisons of primary prostate cancer were performed with cBioPortal using TCGA-PRAD PanCancer dataset. Only samples with mRNA (RNA Seq V2 SEM) and protein (RPPA) expression data were selected for analysis. Samples were grouped based on quartiles of *PLEKHS1* mRNA expression (RNA Seq V2 SEM) or p-SRC Y419 levels (RPPA) and compared for Akt and p-Akt (S437, T308) protein levels (RPPA).

Statistical tests were performed using GraphPad Prism and are described in figure legends. Boxplots were created using R (ggplot2).

### QUANTIFICATION AND STATISTICAL ANALYSIS

#### General statistical analyses.

For pairwise comparisons, two-sided unpaired or paired (depending on experimental design) Student’s *t*-tests with Welch correction were used. When multiple comparisons were performed on a given dataset, p-values were adjusted with FDR correction. In the particular context of the p85^Avi/Avi^ interactome analysis, where the authors wanted to have a more stringent approach to the selection of interactors, a Holm correction was applied.

When data were baseline-corrected prior to statistical analysis (e.g. Extended Data 6: % of *Pten^-/-^)*: multiple two-tailed one-sample *t*-tests were performed (with Holm-Šídák correction).

When more than 2 conditions were compared and two factors were included in the analyses, one-way and 2-way analyses, respectively, were performed followed by Holm-Šídák’s multiple comparisons tests.

Where data showed departure from the assumptions for parametric tests, in particular from normality, a transformation was applied (log or sqrt, depending of the departure) prior to the analysis. Significance was determined as p<0.05.

Statistical analyses were performed using GraphPad Prism 9 and R version 4.1.3.

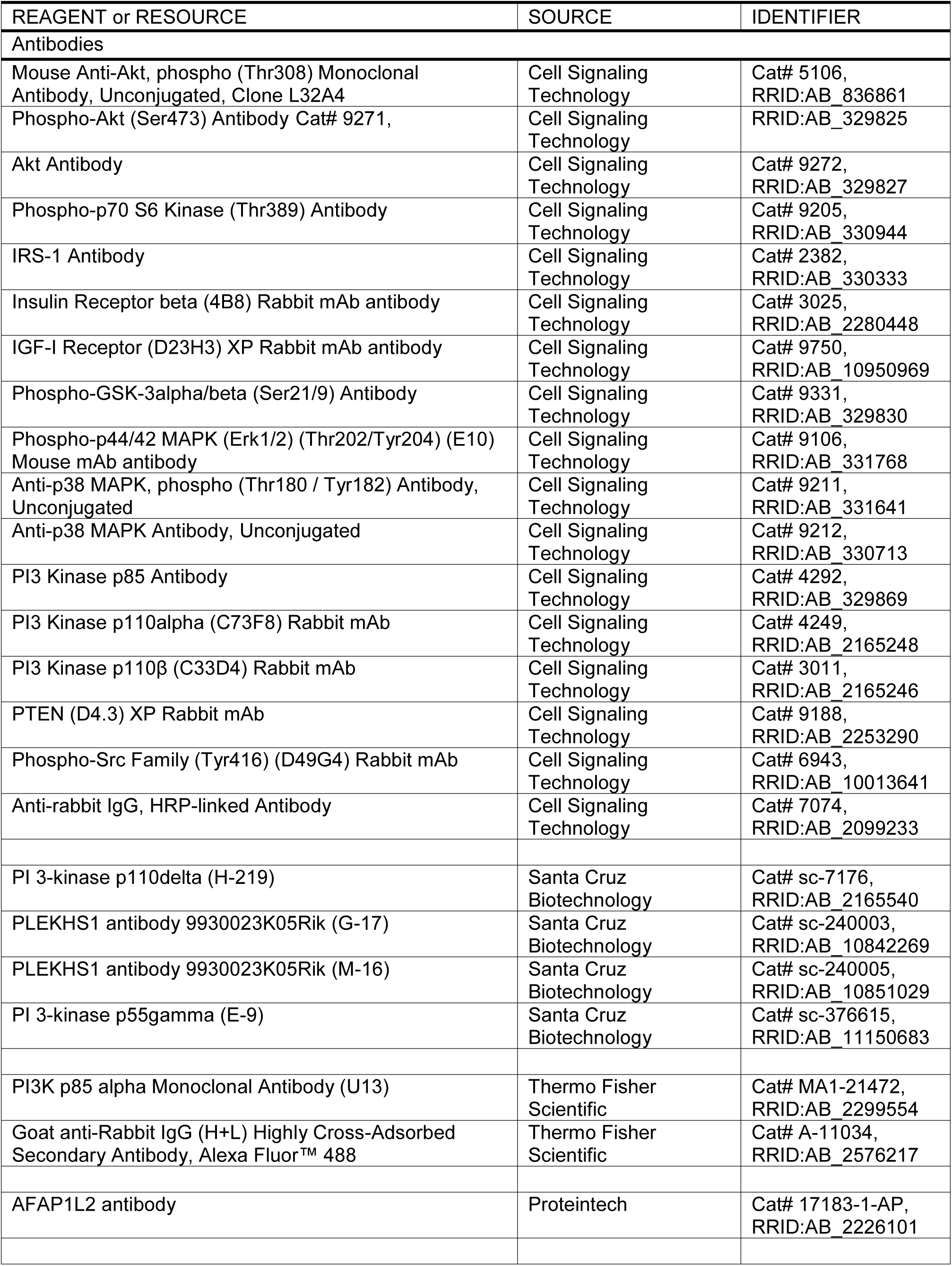
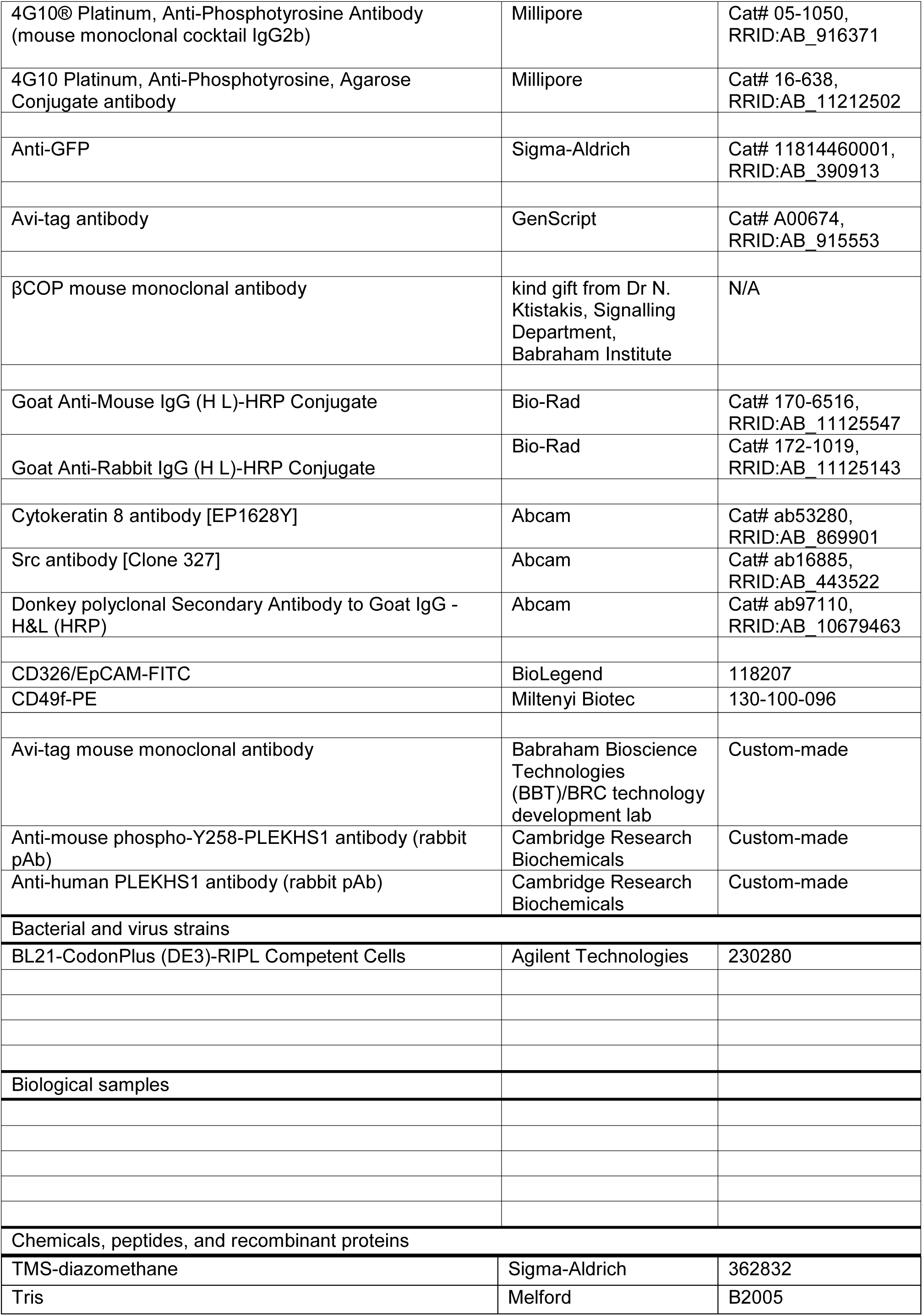
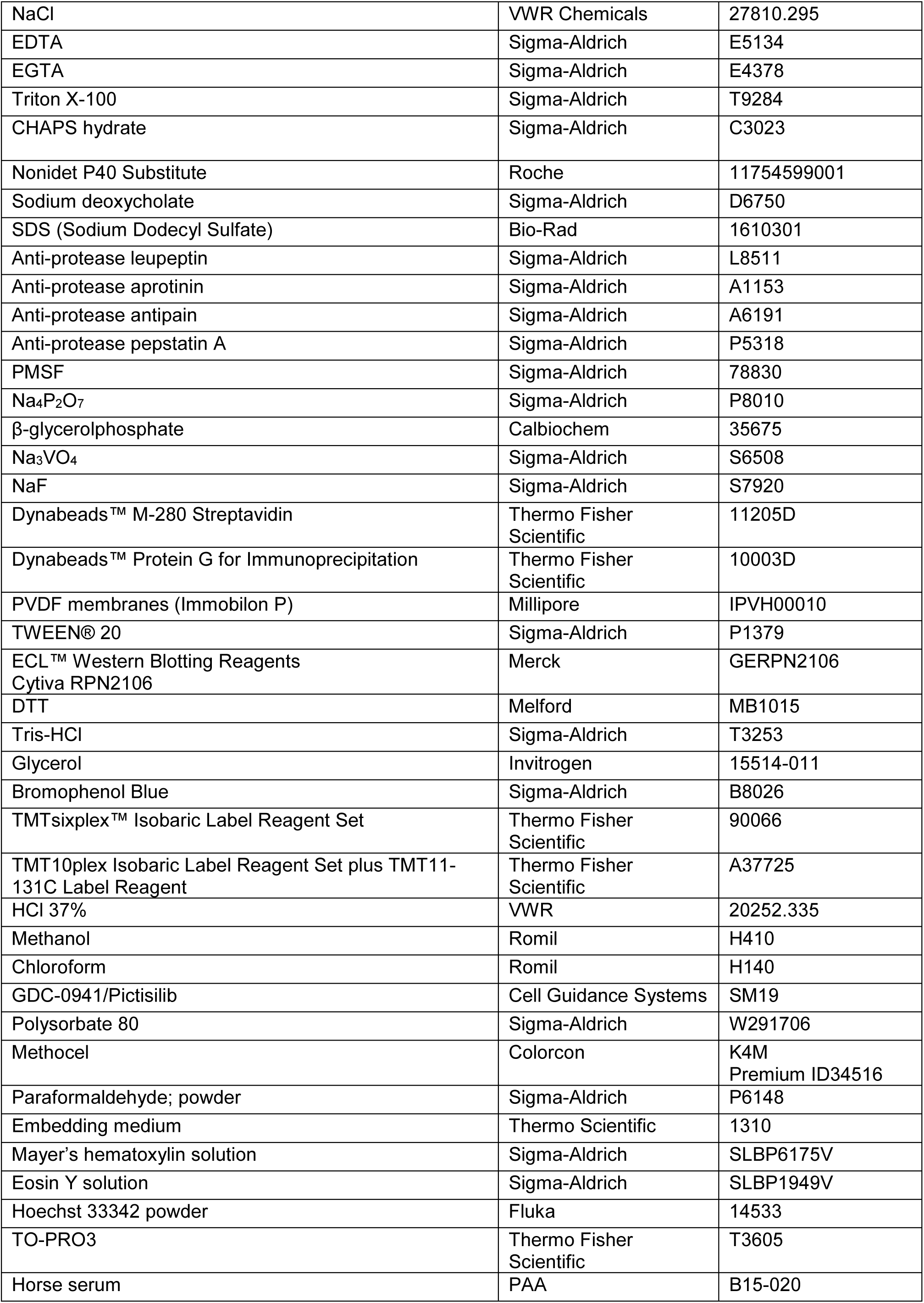
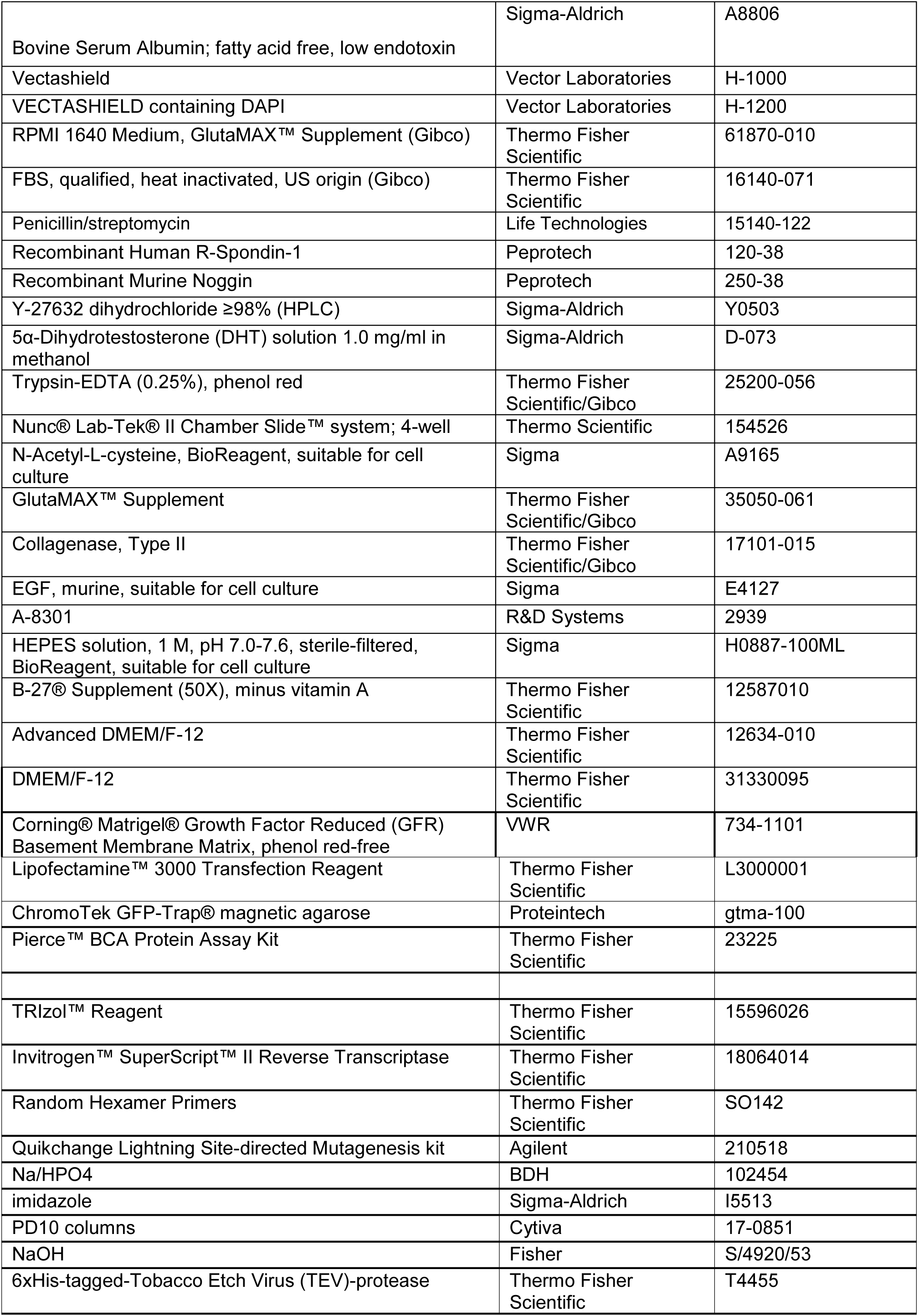
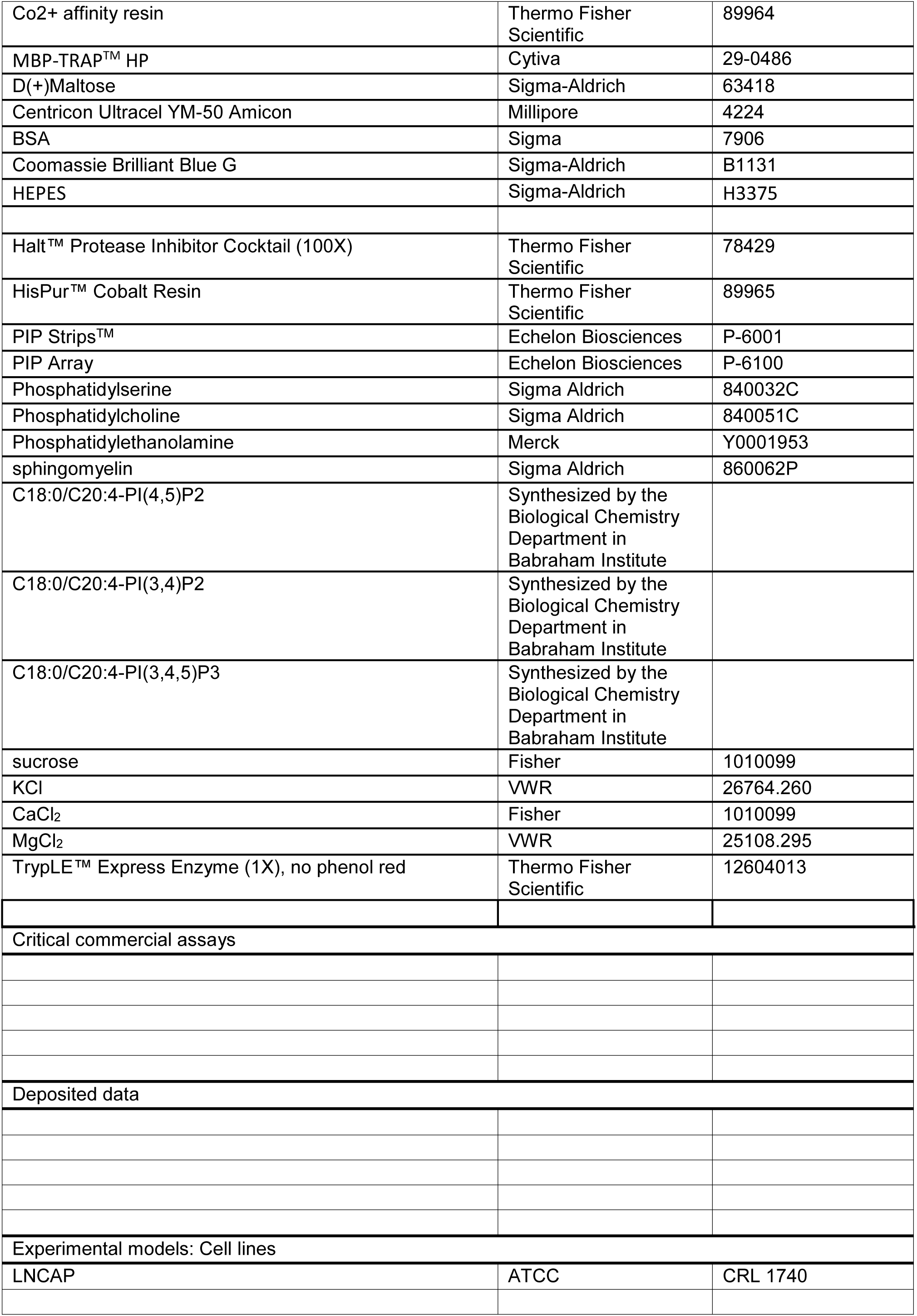
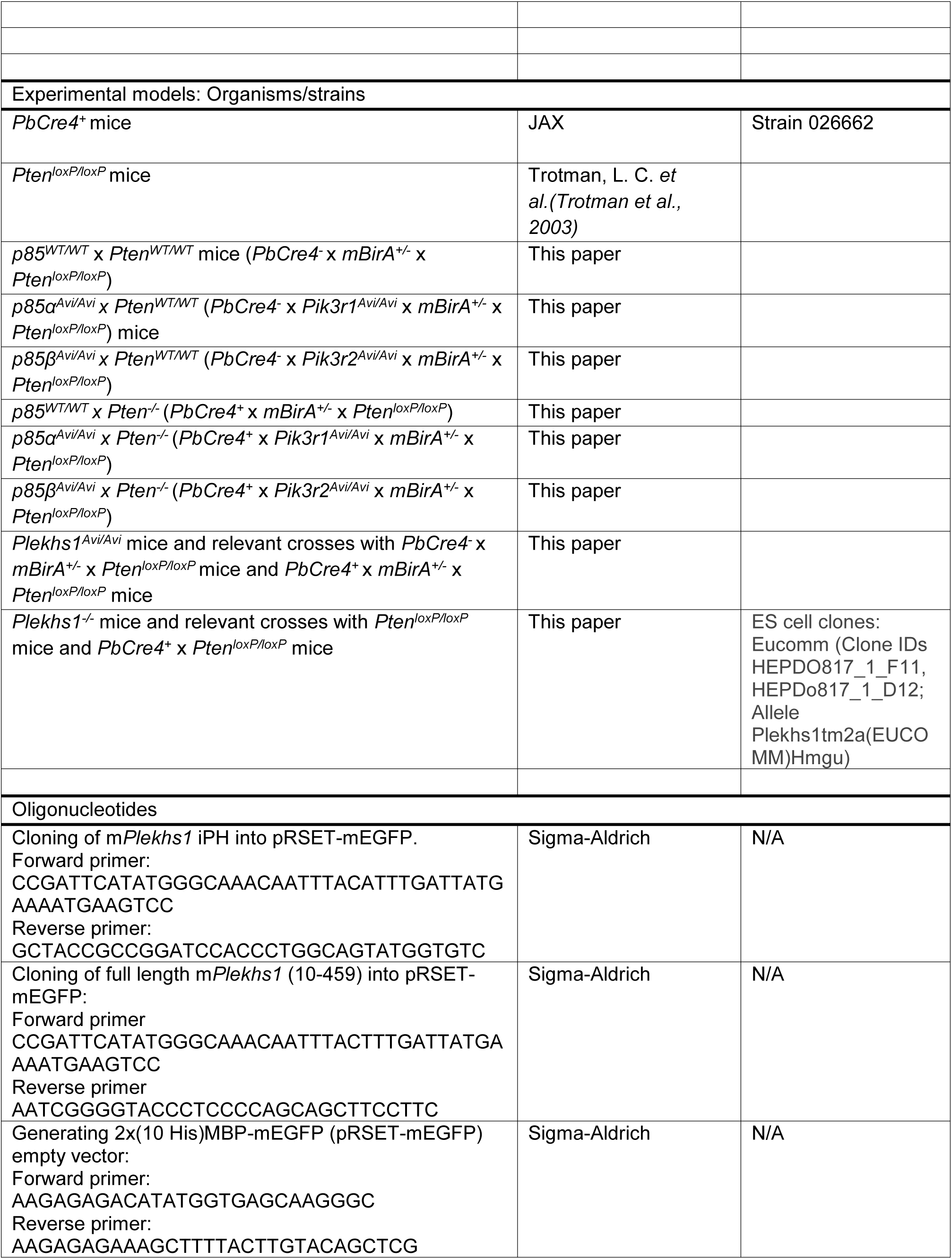
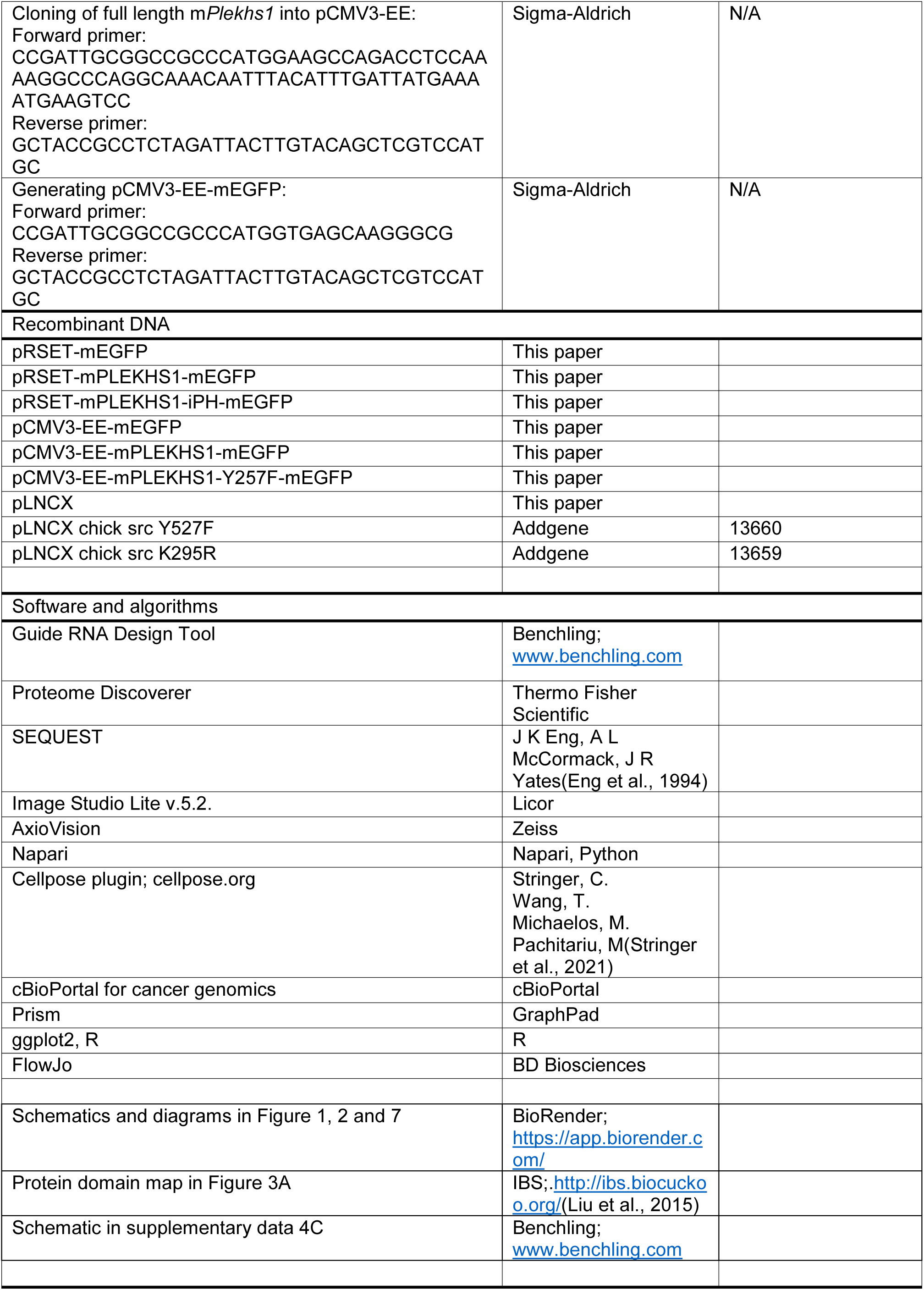
KEY RESOURCES TABLE

