## Supplementary material for "PLEKHS1 drives PI3Ks and remodels pathway homeostasis in PTEN-null prostate": Chessa et al Supplementary Information

### Figure S1

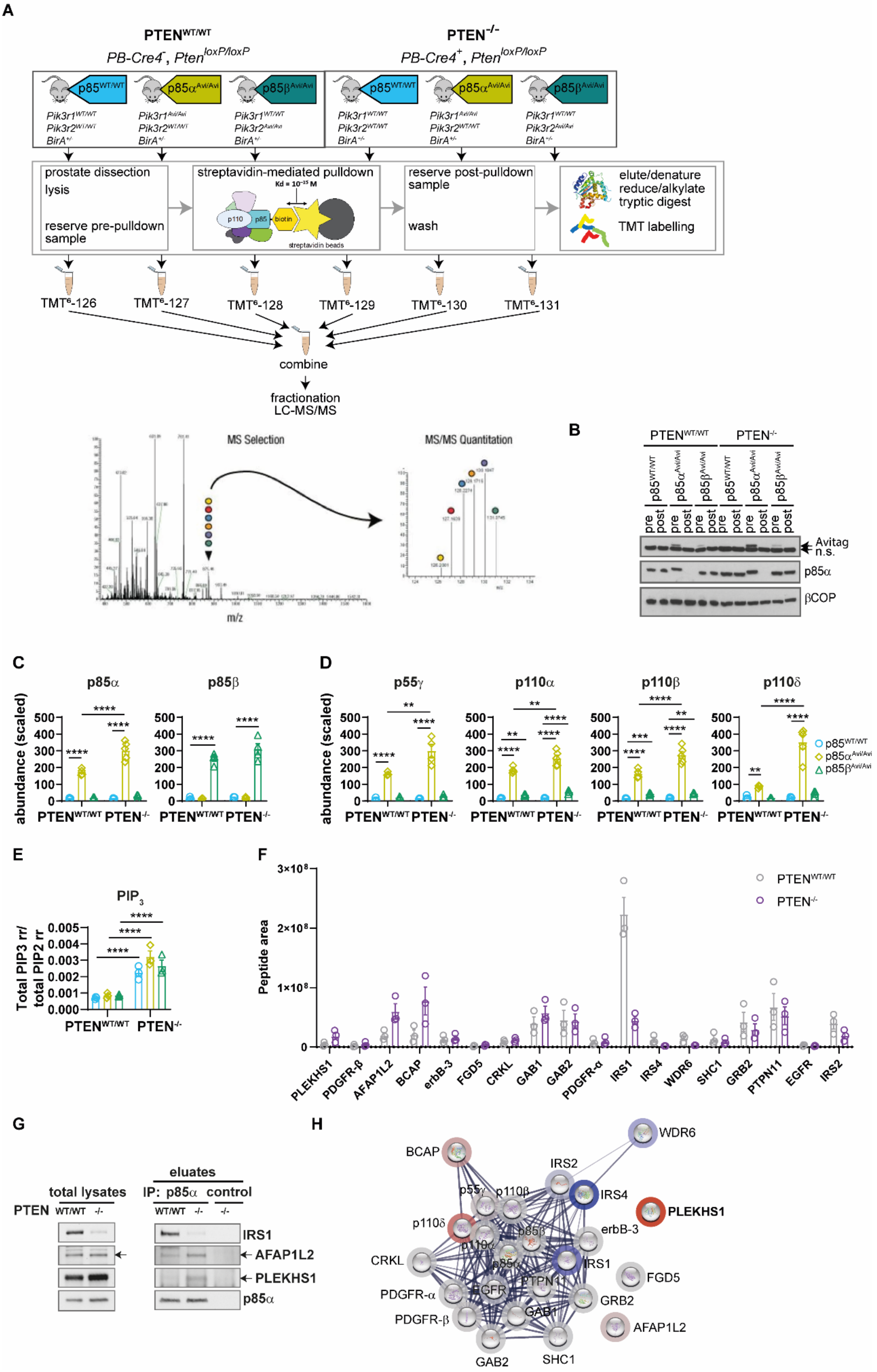

A
PTEN WT/WT
PB-Cre4-, Pten loxP/loxP
PTEN -/-
PB-Cre4+, Pten loxP/loxP
p85 WT/WT
p85α Avi/Avi
p85β Avi/Avi
prostate dissection
lysis
reserve pre-pulldown sample
streptavidin-mediated pulldown
reserve post-pulldown sample
wash
elute/denature
reduce/alkylate
tryptic digest
TMT labelling
TMT6-126
TMT6-127
TMT6-128
TMT6-129
TMT6-130
TMT6-131
combine
fractionation
LC-MS/MS
MS Selection
MS/MS Quantitation
m/z
B
Avitag
n.s.
p85α
βCOP
C
p85α
p85β
abundance (scaled)
D
p55γ
p110α
p110β
p110δ
E
PIP3
Total PIP3 rr/ total PIP2 rr
F
Peptide area
PLEKHS1
PDGFR-β
AFAP1L2
BCAP
erbB-3
FGD5
CRKL
GAB1
GAB2
PDGFR-α
IRS1
IRS4
WDR6
SHC1
GRB2
PTPN11
EGFR
IRS2
G
total lysates
eluates
IP: p85α control
IRS1
AFAP1L2
PLEKHS1
p85α
H
WDR6
BCAP
IRS2
IRS4
PLEKHS1
erbB-3
CRKL
FGD5
PDGFR-α
GRB2
PDGFR-β
AFAP1L2
GAB2
SHC1

### Figure S2

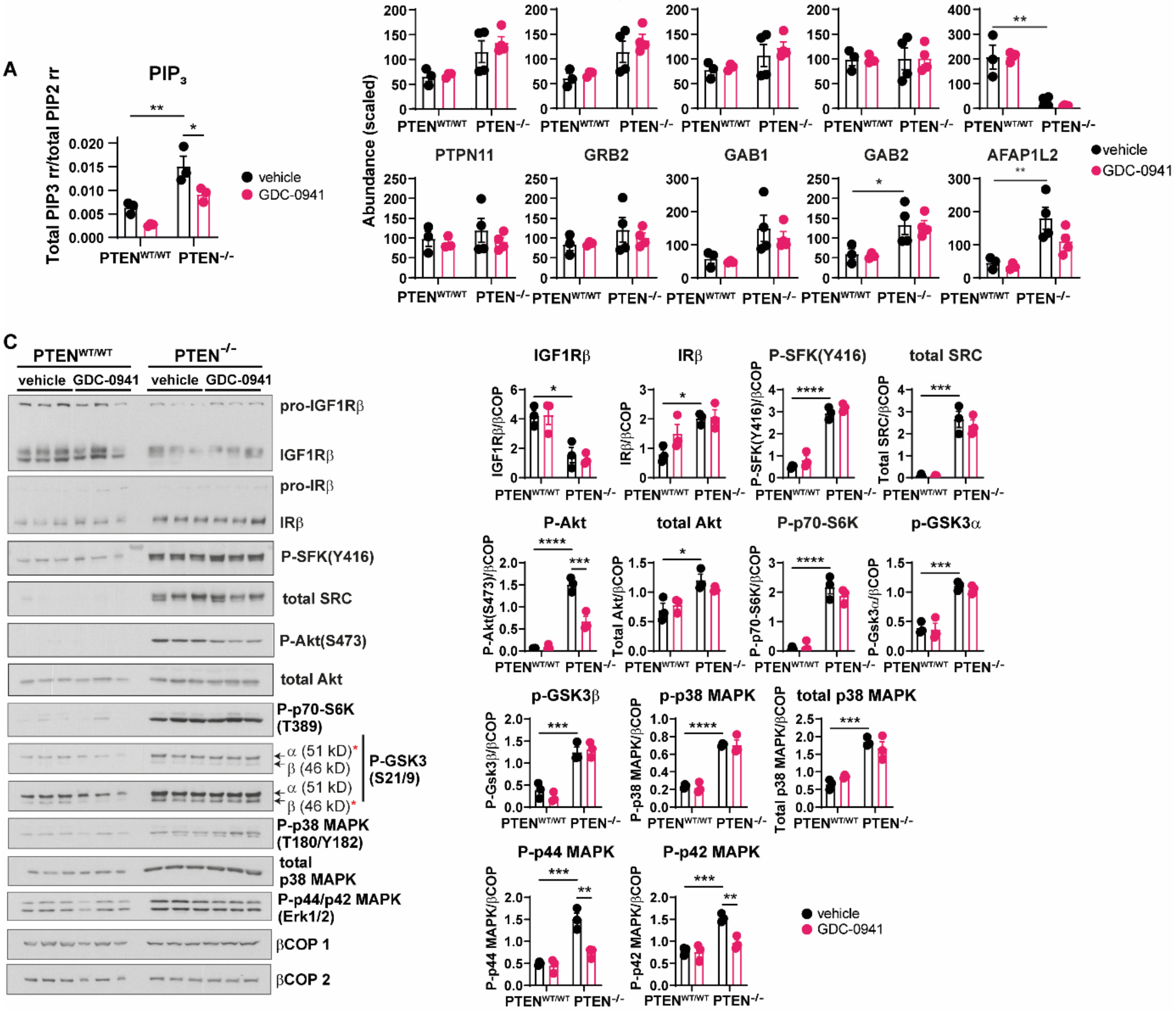
A
PIP3
Total PIP3 rr/total PIP2 rr
vehicle
GDC-0941
PTENWT/WT PTEN-/-
B
Abundance (scaled)
p85α p55γ p110α p110β IRS4
PTPN11 GRB2 GAB1 GAB2 AFAP1L2
C
PTENWT/WT PTEN-/-
vehicle GDC-0941 vehicle GDC-0941
pro-IGF1Rβ
IGF1Rβ
pro-IRβ
IRβ
P-SFK(Y416)
total SRC
P-Akt(S473)
total Akt
P-p70-S6K (T389)
α (51 kD)
β (46 kD)
P-GSK3 (S21/9)
P-p38 MAPK (T180/Y182)
total p38 MAPK
P-p44/p42 MAPK (Erk1/2)
βCOP 1
βCOP 2
IGF1Rβ IRβ P-SFK(Y416) total SRC
P-Akt total Akt P-p70-S6K p-GSK3α
p-GSK3β p-p38 MAPK total p38 MAPK
P-p44 MAPK P-p42 MAPK

**Figure S3**

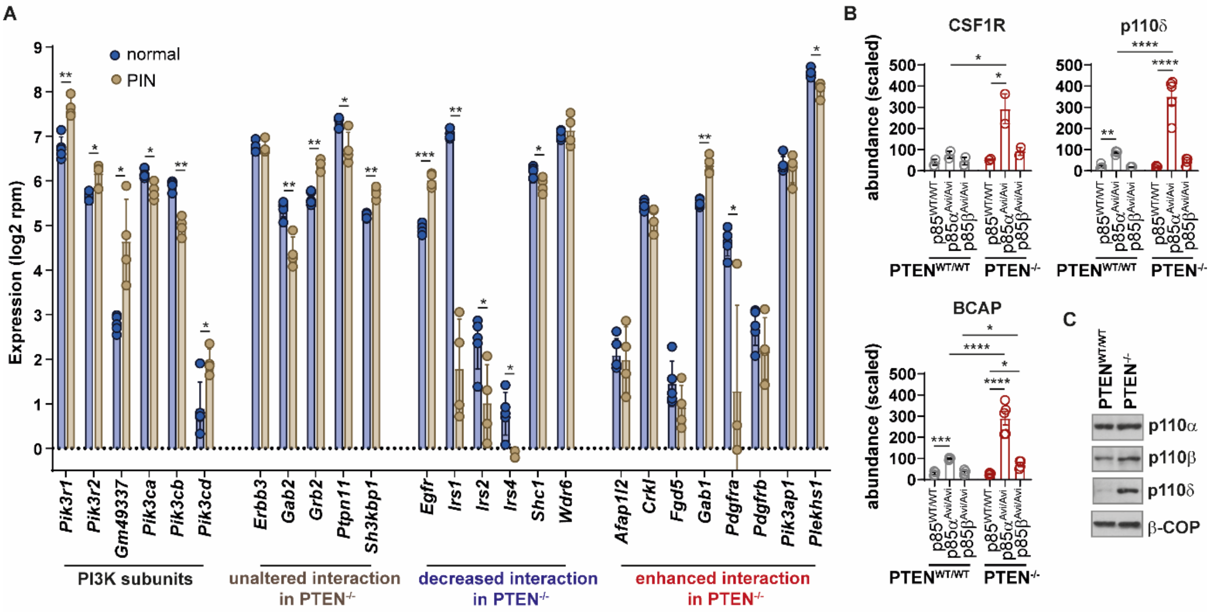
A
normal
PIN
Expression (log2 rpm)
Pik3r1
Pik3r2
Gm49337
Pik3ca
Pik3cb
Pik3cd
Erbb3
Gab2
Grb2
Ptpn11
Sh3kbp1
Egfr
Irs1
Irs2
Irs4
Shc1
Wdr6
Afap1l2
Crkl
Fgd5
Gab1
Pdgfra
Pdgfrb
Pik3ap1
Plekhs1
PI3K subunits
unaltered interaction in PTEN-/-
decreased interaction in PTEN-/-
enhanced interaction in PTEN-/-
B
CSF1R
p110δ
BCAP
abundance (scaled)
PTENWT/WT
PTEN-/-
C
p110α
p110β
p110δ
β-COP

**Figure S4**

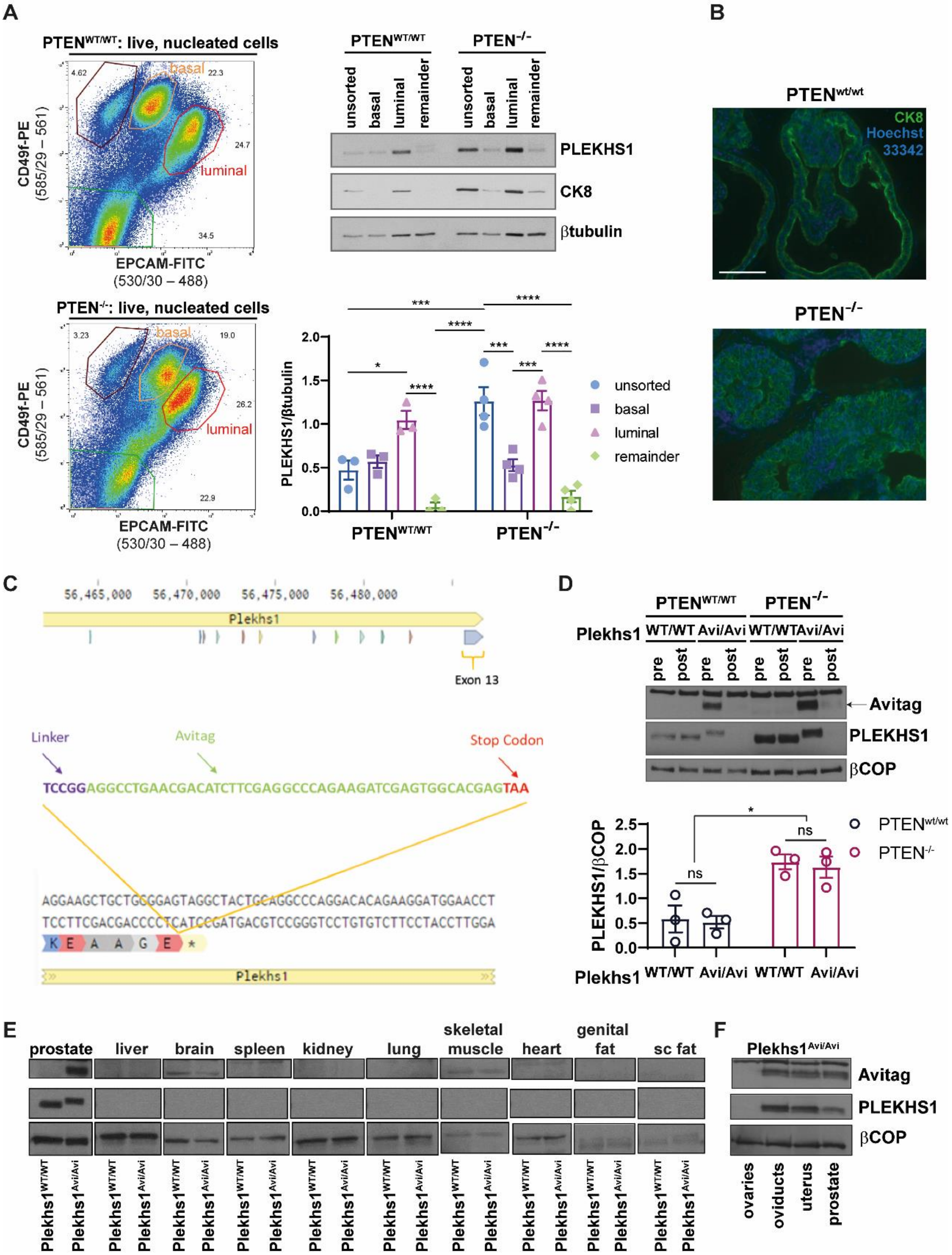
A
B
PTEN^WT/WT: live, nucleated cells
PTEN-/-: live, nucleated cells
basal
luminal
CD49f-PE
(585/29 – 561)
EPCAM-FITC
(530/30 – 488)
unsorted
basal
luminal
remainder
PLEKHS1
CK8
βtubulin
PLEKHS1/βtubulin
PTENwt/wt
PTEN-/-
CK8
Hoechst
33342
C
Plekhs1
Exon 13
Linker
Avitag
Stop Codon
TCCGGAGGCCTGAACGACATCTTCGAGGCCCAGAAGATCGAGTGGCACGAGTAA
D
Plekhs1
WT/WT
Avi/Avi
pre
post
Avitag
PLEKHS1
βCOP
PLEKHS1/βCOP
ns
E
prostate
liver
brain
spleen
kidney
lung
skeletal muscle
heart
genital fat
sc fat
F
Plekhs1Avi/Avi
ovaries
oviducts
uterus
prostate

Figure S5

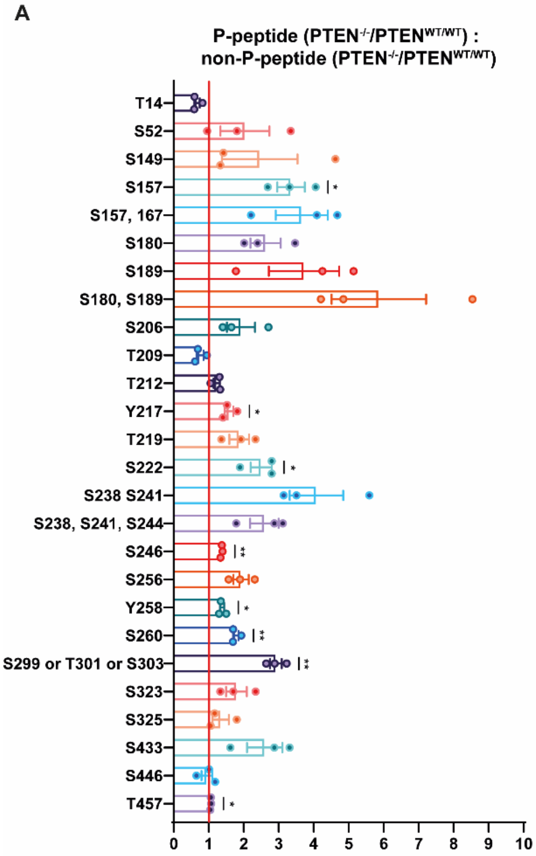
A
P-peptide (PTEN-/-/PTENWT/WT) :
non-P-peptide (PTEN-/-/PTENWT/WT)
T14
S52
S149
S157
S157, 167
S180
S189
S180, S189
S206
T209
T212
Y217
T219
S222
S238 S241
S238, S241, S244
S246
S256
Y258
S260
S299 or T301 or S303
S323
S325
S433
S446
T457
0 1 2 3 4 5 6 7 8 9 10

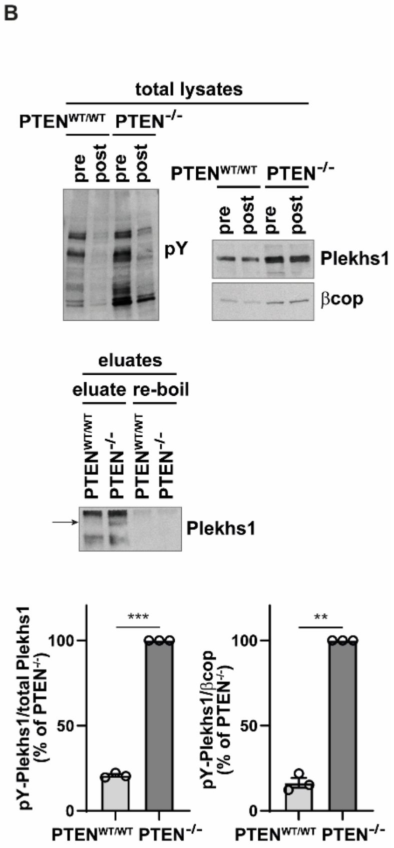
B
total lysates
PTENWT/WT PTEN-/-
pre post pre post
pY
PTENWT/WT PTEN-/-
pre post pre post
Plekhs1
βcop
eluates
eluate re-boil
PTENWT/WT PTEN-/- PTENWT/WT PTEN-/-
Plekhs1
pY-Plekhs1/total Plekhs1 (% of PTEN-/-)
***
100
50
0
PTENWT/WT PTEN-/-
pY-Plekhs1/βcop (% of PTEN-/-)
**
100
50
0
PTENWT/WT PTEN-/-

### Figure S6

**A**

| | ***Plekhs1***$^{WT/WT}$ | ***Plekhs1***$^{+/-}$ | ***Plekhs1***$^{-/-}$ |
|---|---|---|---|
| **Observed** | 66 | 111 | 48 |
| **Expected** | 72 | 145 | 72 |

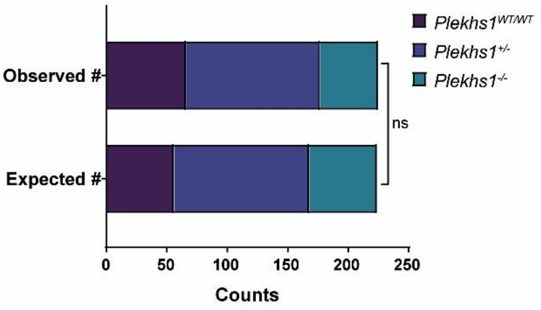


**B**

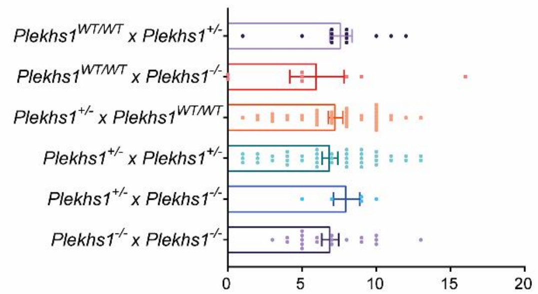


**C**

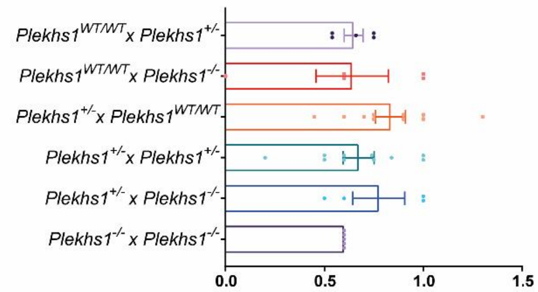


**D**

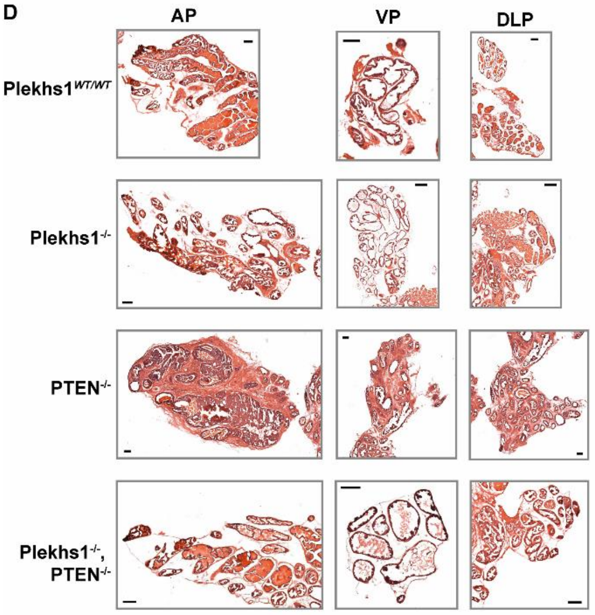


**E**

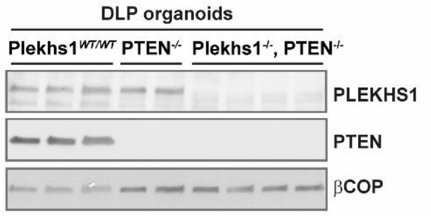


**F**

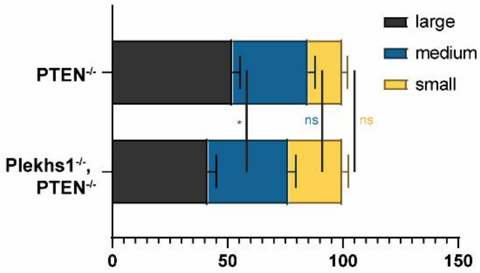

**Figure S7**

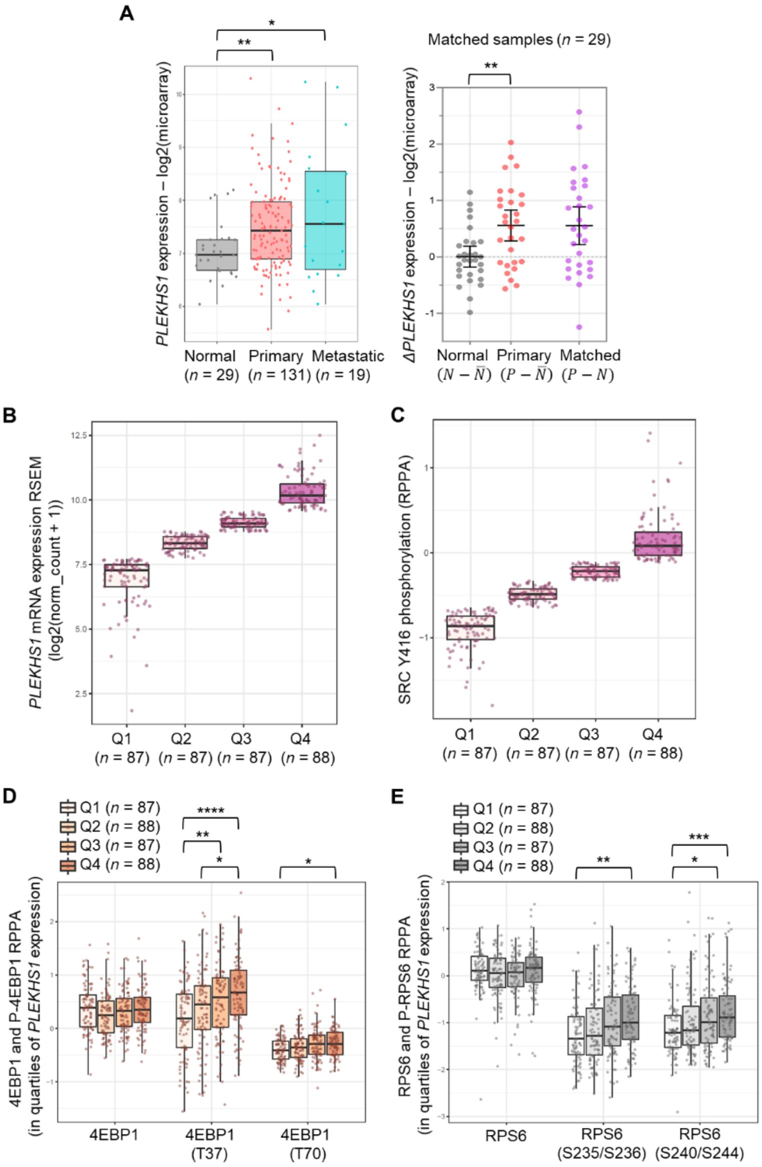
A
PLEKHS1 expression – log2(microarray)
Normal (n = 29)
Primary (n = 131)
Metastatic (n = 19)
Matched samples (n = 29)
ΔPLEKHS1 expression – log2(microarray)
Normal (N − N̄)
Primary (P − N̄)
Matched (P − N)
B
PLEKHS1 mRNA expression RSEM (log2(norm_count + 1))
Q1 (n = 87) Q2 (n = 87) Q3 (n = 87) Q4 (n = 88)
C
SRC Y416 phosphorylation (RPPA)
Q1 (n = 87) Q2 (n = 87) Q3 (n = 87) Q4 (n = 88)
D
Q1 (n = 87)
Q2 (n = 88)
Q3 (n = 87)
Q4 (n = 88)
4EBP1 and P-4EBP1 RPPA (in quartiles of PLEKHS1 expression)
4EBP1
4EBP1 (T37)
4EBP1 (T70)
E
Q1 (n = 87)
Q2 (n = 88)
Q3 (n = 87)
Q4 (n = 88)
RPS6 and P-RPS6 RPPA (in quartiles of PLEKHS1 expression)
RPS6
RPS6 (S235/S236)
RPS6 (S240/S244)

**Figure S1. Endogenous class 1A PI3K subunits are efficiently recovered by streptavidin mediated pulldown of p85α-Avi and p85β-Avi, related to Figure 1.** **A,** Full workflow for identification of p85α- and p85β-interactors in 12-week-old *Pten*$^{WT/WT}$ and *Pten*$^{-/-}$ prostates. Genotypes of mice/prostates used for pulldown of Avi-tagged, endogenous p85α and p85β are indicated. Prostates were rapidly dissected, flash-frozen in liquid nitrogen and lysed. Lysates were subjected to streptavidin-mediated pulldown and proteins eluted, purified by SDS-PAGE and digested in gel. TMT-6-Plex labelling of peptides was performed according to manufacturer's instructions. Samples were combined, fractionated and subjected to TMT-LC-MS/MS. Quantification of proteins and calculation of scaled abundances was performed in Proteome Discoverer. **B,** Immuno-blots of pre-and post-pulldown samples, probed with the indicated antibodies showing; efficient depletion of Avi-tagged proteins in post-pulldown lanes, the small increase in size of the Avi-tagged p85s and that there is about 5x more p85α than p85β in prostate tissue. The non-specific band, decorated with the Avi-tag Genscript antibody, is indicated as n.s. A representative blot of n=4 independent experiments is shown. An estimated total of 20 µg protein was loaded per lane. **C, D,** Scaled abundances (each interactor is given a total of 3200 abundance units that are scaled across the 6 different conditions and replicates) of PI3K regulatory subunits (**C**) and catalytic subunits (**D**) detected in p85α- and p85β-Avi pulldowns run as 4 cohorts of n=4-5 separate biological replicates per genotype (n=4 biological replicates for all genotypes, except *p85α*$^{Avi/Avi}$ x *Pten*$^{-/-}$ where n=5 (here, n4 and n5 were in same experimental cohort and average of 2 technical replicates)). Data are means ± SEM. For statistical analysis, a 2-way ANOVA was performed on sqrt transformed data followed by Holm-Šídák's multiple comparisons tests. Adjusted P value summaries <0.05 are indicated. **E,** $PIP_3$ measurements in prostates of the indicated genotypes. Data are means ± SEM of 3 biological replicates per genotype and are expressed as the ratio of the abundance of total $PIP_3$ to that of total $PIP_2$ in the same sample. For statistical analysis, a 2-way ANOVA was performed on log transformed data followed by Holm-Šídák's multiple comparisons tests. **F,** Peptide areas (sum of the 3 most abundant peptides for each protein, which is thus a measure of the relative moles of each protein) as calculated in Proteome Discoverer software. Data are means ± SEM of n=3 biological replicates per genotype, run in separate cohorts. **G,** IP from PtenWT/WT and Pten-/- prostates (12 weeks) using p85α or control antibody (βCOP). Total lysates (20 µg/lane) and IP eluates (2 mg total protein input) were blotted with the indicated antibodies. A representative blot from 3 independent experiments is shown. When extracted from mouse tissues and cells, mouse (and human) PLEKHS1 migrates with an estimated relative mass of about 70kD during SDS-PAGE, although the longest predicted open reading frame in mouse has a predicted relative mass of 53kD. We have observed that bacterially-expressed, tagged-versions of mPLEKHS1 also migrate with an estimated relative mass of 70kD during SDS-PAGE (correcting for the size of the tags). We assume this is an intrinsic property of PLEKHS1 peptide sequence and may be a result of its low isoelectric point (6.05) and predicted charge of -23 at pH 8.0, leading to reduced SDS-binding and migration speed during PAGE. **H,** STRING network analysis of the p85α-Avi and p85β-Avi interactome in *Pten*$^{WT/WT}$ and *Pten*$^{-/-}$ prostates (12 weeks) (https://string-db.org/), using the Proteins with Values/Ranks functionality, where input values are log2 fold change (FC) between *Pten*$^{-/-}$:*Pten*$^{WT/WT}$. The halo colour is based on the rank of the protein in the full, sorted set of input values. Network nodes represent proteins, lines represent known protein-protein associations (functional and physical interactions). The thickness of the line corresponds to the confidence of the protein-protein association.

**Figure S2. Effect of pan-PI3K inhibition on growth factor signalling and PI3K interactions, related to Figure 2. A,** $PIP_3$ measurements in *p85α$^{Avi/Avi}$x Pten$^{WT/WT}$* and *p85α$^{Avi/Avi}$x Pten$^{-/-}$* prostates (24-32 weeks), treated with vehicle or GDC-0941. Data are means ± SEM of 3 biological replicates per genotype/condition and are expressed as the ratio of the abundance of total $PIP_3$ to that of total $PIP_2$ in the same sample. For statistical analysis, a 2-way ANOVA was performed followed by Holm-Šídák's multiple comparisons tests. Significant p value summaries (<0.05) are indicated. **B,** Targeted TMT-LC-MS/MS analysis of the indicated proteins in *p85α$^{Avi/Avi}$x Pten$^{WT/WT}$* and *p85α$^{Avi/Avi}$x Pten$^{-/-}$* prostates (24-32 weeks), treated with vehicle or GDC-0941 (scaled abundances, where each interactor is given a total of 1400 abundance units that are scaled across the different conditions and replicates). Data are means ± SEM for n=3-4 biological replicates per genotype/condition, run in 2 separate cohorts (*p85α$^{Avi/Avi}$x Pten$^{WT/WT}$*: n=3; *p85α$^{Avi/Avi}$, Pten$^{-/-}$:* n=4). For statistical analysis, a 2-way ANOVA was performed followed by Holm-Šídák's multiple comparisons tests. Significant p value summaries (<0.05) are indicated. **C**: Immunoblots with the indicated antibodies of total lysates from *p85α$^{Avi/Avi}$x Pten$^{WT/WT}$* and *p85α$^{Avi/Avi}$x Pten$^{-/-}$* prostates (24-32 weeks), treated with vehicle or GDC-0941 (20 µg total protein/lane). Data are normalised to βCOP1; except IRβ and P-GSK3α/β, which are normalised to βCOP2. Data are means ± SEM of 3 biological replicates per genotype/condition, run as a single cohort. For statistical analysis, a 2-way ANOVA was performed followed by Holm-Šídák's multiple comparisons tests. Significant p value summaries (<0.05) are indicated.

**Figure S3. Expression levels of p85 interacting proteins, and p85 interactions with immune cell components, related to Figure 1 and S1. A,** RNAseq data from *Pten$^{WT/WT}$* and *Pten$^{-/-}$* anterior prostate lobes (NCBI Gene Expression Omnibus accession GSE94574) at the PIN stage of tumourigenesis; re-analysed and quantitated in SeqMonk and expressed as log2RPM values. For statistical analysis, multiple unpaired *t*-tests were performed (Welch correction); a FDR correction was applied for multiple comparisons (two-stage step-up (Benjamini, Krieger, and Yekutieli)). Q value summaries <0.05 are shown. **B**, Scaled abundances of indicated proteins identified as p85α-Avi and/or p85β-Avi interactors in prostate (12 weeks). Data are mean ± SEM of n=4-5 biological replicates per genotype (n=4 for all genotypes, except p85α$^{Avi/Avi}$, PTEN$^{-/-}$ where n=5), except for CSF1R, a low abundance protein detected in n=2 biological replicates, where data are mean ± range. For statistical analysis, a 2-way ANOVA was performed followed by Holm-Šídák's multiple comparisons tests. Adjusted P value summaries <0.05 are indicated. **C,** Total lysates from *Pten$^{WT/WT}$* and *Pten$^{-/-}$* prostates (12 weeks, 25 µg/lane) immunoblotted with the indicated antibodies. A representative blot from 2 independent experiments is shown.

**Figure S4. PLEKHS1 expression is tissue-restricted, enhanced in *Pten*$^{-/-}$ whole prostate and enriched in luminal epithelial cells, related to Figure S1 and 4. A,** Immunoblots of FACS sorted prostate cell populations. Left: gating strategy, showing percentages of basal and luminal epithelial cells, as well as EPCAM-negative populations (remainder). Right: a total of 120000 cells were loaded per lane for each cell population. A representative blot from n=3 independent experiments is shown. CK8 is a luminal cell marker. PLEKHS1 expression (G17 antibody) was quantified and normalised to βtubulin. Data are means ± SEM of n=3 (*Pten*$^{WT/WT}$ populations) or n=4 (*Pten*$^{-/-}$ populations) biological replicates. For statistical analysis, a 2-way ANOVA was performed followed by Holm-Šídák's multiple comparisons tests. Significant p value summaries (<0.05) are indicated. **B,** Widefield fluorescence images of *Pten*$^{WT/WT}$ and *Pten*$^{-/-}$ prostate cryosections (12 weeks) stained with the indicated antibodies/dyes. Scalebar: 100 µm. **C,** Schematic of Plekhs1$^{Avi/Avi}$ gene targeting strategy. **D,** Prostate lysates of the indicated genotypes (30 µg/lane) were immunoblotted with Avitag (Genscript) or PLEKHS1 (G17). PLEKHS1 expression was quantified and normalised to βCOP. Data are means ± SEM of n=3 biological replicates per genotype, run as 3 independent experiments. For statistical analysis, a 2-way ANOVA was performed followed by Holm-Šídák's multiple comparisons tests. **E,** *Plekhs1*$^{WT/WT}$*x Pten*$^{WT/WT}$ and *Plekhs1*$^{Avi/Avi}$*x Pten*$^{WT/WT}$ lysates from the indicated tissues were immunoblotted with Avitag (Genscript), PLEKHS1 G17 and βCOP. A total of 25 µg protein was loaded per lane, with the exception of subcutaneous (sc) fat and genital fat, where 12.5 µg was loaded. **F,** *Plekhs1*$^{Avi/Avi}$*x Pten*$^{WT/WT}$ lysates from the indicated tissues were immunoblotted as described in e. A total of 30 µg protein was loaded per lane.

**Figure S5. PLEKHS1 phospho-site analysis, related to Figure 4 and 5. A,** Quantitation of phospho-peptides identified in Plekhs1-Avi pulldown and TMT-LC-MS/MS. Data are ratios of the signals for the phosphopeptide, normalised to the equivalent non-phospho-peptide, in *Plekhs1*$^{Avi/Avi}$*x Pten*$^{-/-}$ *: Plekhs1*$^{Avi/Avi}$*x Pten*$^{WT/WT}$. Data are mean ± SEM of n=3 biological replicates per genotype. For statistical analysis, two-tailed one-sample *t*-tests were performed (with Holm-Šídák correction; comparison of each *Plekhs1*$^{Avi/Avi}$*x Pten*$^{-/-}$ *: Plekhs1*$^{Avi/Avi}$*x Pten*$^{WT/WT}$ ratio to hypothetical mean of 1 (*Plekhs1*$^{Avi/Avi}$*x Pten*$^{WT/WT}$ *: Plekhs1*$^{Avi/Avi}$*x Pten*$^{WT/WT}$)). Significant p value summaries (<0.05) are indicated. **B,** Phospho-tyrosine IP from PTEN$^{WT/WT}$ and PTEN$^{-/-}$ prostates from mice (12 weeks), immuno-blotted with the indicated antibodies. For PLEKHS1, the G17 antibody was used. A representative blot of total lysates (25 µg total protein per lane) and IP eluates (from 2 mg total protein input) is shown. PLEKHS1 signal in the IP eluates was quantified as pY-PLEKHS1 and normalised to total PLEKHS1 or βCOP signal in total lysates. Data are means ± SEM of n=3 biological replicates per genotype, run as 3 independent experiments. For statistical analysis, multiple two-tailed one-sample *t*-tests were performed (with Holm-Šídák correction) on baseline-corrected data (% of *Pten*$^{-/-}$*).* P value summaries are indicated.

**Figure S6. Genetic ablation of Plekhs1 does not adversely affect healthy prostate development, however, *Pten*$^{-/-}$ prostate tumour development is delayed and the proportion of large organoids is significantly reduced, related**

**to Figure 6. A**, Top: the numbers of *Plekhs1$^{WT/WT}$, Plekhs1$^{+/-}$* and *Plekhs1$^{-/-}$* mice born from *Plekhs1$^{+/-}$ x Plekhs1$^{+/-}$* crosses. The expected number of mice was based on the expected Mendelian 1:2:1 ratio, calculated according to the total number of mice born. Bottom: Chi square analysis to determine whether there was a significant difference from expected ratio. p value summary is shown. **B, C,** litter size (**B**) and litter frequency (**C**) of mice born from the indicated crosses. Data are means ± SEM of n=42 (*Plekhs1$^{+/-}$ x Plekhs1$^{+/-}$*), n=39 (*Plekhs1$^{+/-}$ x Plekhs1$^{WT/WT}$*), n=20 (*Plekhs1$^{-/-}$ x Plekhs1$^{-/-}$*), n=13 (*Plekhs1$^{WT/WT}$ x Plekhs1$^{+/-}$*), n=8 (*Plekhs1$^{WT/WT}$ x Plekhs1$^{-/-}$*) and n=5 (*Plekhs1$^{+/-}$ x Plekhs1$^{-/-}$*). For statistical analysis, an ordinary one-way ANOVA was performed (Holm-Šídák's multiple comparisons test). None of the comparisons in B and C were significant. **D,** H&E-stained cryosections of prostates taken from the indicated lobes and genotypes (12-15 weeks). Images were acquired using a Zeiss MicroBeam microscope, and automatically stitched with AxioVision software, with shading correction enabled. AP: anterior prostate; VP: ventral prostate; DLP: dorsolateral prostate. Representative images from n=3 biological replicates are shown. Scalebars: 200 µm. **E,** Total lysates from *Pten$^{WT/WT}$, Pten$^{-/-}$* and *Plekhs1$^{-/-}$x Pten$^{-/-}$* DLP organoids (derived from 12-15 week old mice) at passage 0, day 11 of culture, were immuno-blotted with the indicated antibodies. A total of 15 µg protein was loaded per lane. **F,** Average organoid area, categorised by size (small, medium and large), in *Pten$^{-/-}$* and *Plekhs1$^{-/-}$*, *Pten$^{-/-}$* DLP organoids (derived from 13-14 week old mice), at passage 0, day 6 of culture. Organoid area was measured as described in Figure 5I. Data are mean ± SD of n=7 biological replicates per genotype. For statistical analysis, a 2-way ANOVA was performed (Holm-Šídák's multiple comparisons test). P value summaries are indicated.

**Figure S7. *PLEKHS1* mRNA expression is increased in prostate cancer and correlates with PI3K/Akt/mTOR pathway activation, related to Figure 7. A,** PLEKHS1 gene expression (GPL10264 Affymetrix microarray, probe ID 17458, NM_024889 transcript) in Taylor et al. (2010) dataset. Left: Pooled analysis of PLEKHS1 expression shows significant increase in primary (p=0.00459, n=131, LIMMA analysis) and metastatic (p=0.0109, n=19, LIMMA analysis) samples compared to normal prostate (n=29). Right: PLEKHS1 expression is significantly higher in primary tumour samples (P) compared to matched normal benign prostate (N) (p=0.003, n=29, Wilcoxon test). Scatterplot shows mean and 95% confidence intervals. **B-C,** Primary prostate cancer samples were grouped by quartiles of *PLEKHS1* mRNA expression (RNA Seq V2 RSEM) (**B**) or SRC Y416 phosphorylation (RPPA) (**C**). Data from cBioPortal TCGA-PRAD. **D-E,** Primary prostate cancer samples were grouped by quartiles of *PLEKHS1* mRNA expression (RSEM) and compared for 4EBP1, p-4EBP1 (T37, T70), RPS6, p-RPS6 (S235/S236, S240/S244) protein levels (RPPA). Data from cBioPortal TCGA-PRAD. Statistics: Kruskal-Wallis with Dunn’s multiple comparison test.
